# Conformational Changes of the ABC Transporter BmrA Depend on Membrane Curvature

**DOI:** 10.1101/2024.01.24.577054

**Authors:** Alicia Damm, Kemil Belhadji, Raj Kumar Sadhu, Su-Jin Paik, Aurélie Di-Cicco, John Manzi, Michele Castellana, Raju Regmi, Emmanuel Margeat, Maxime Dahan, Pierre Sens, Daniel Lévy, Patricia Bassereau

## Abstract

While mechanosensitive ion channels’ gating has been well documented, the effect of membrane mechanics, in particular membrane curvature, on the function of transporters remains elusive. Since conical shape transmembrane proteins locally deform membranes, conversely membrane bending could impact their conformations and their function. We tested this hypothesis using BmrA, a bacterial ABC exporter that exhibits large conformational changes upon ATP hydrolysis, switching between open and closed states with opposite V-shapes. After reconstitution in liposomes of different curvatures, and at two different temperatures, we showed that BmrA ATPase activity decreases by 2.9-fold when their diameter decreases from 125 to 29 nm. Moreover, using single-molecule FRET, we observed that the fraction of closed conformations is reduced in highly curved vesicles when adding ATP or non-hydrolysable AMP-PNP. Our results are well explained by a theoretical 2-states model including the effect of membrane mechanics on protein shape transition. Our work reveals that the functional cycle of conical transporters is curvature sensitive, to an extent depending on protein geometry.

**Significance Statement:** Biological membranes actively regulate protein function. While some ion channels are known to sense membrane tension, whether transporters also respond to mechanical cues was unknown. We show that the conical-shaped bacterial exporter BmrA is sensitive to membrane curvature, with both its activity and conformation landscape determined by the surrounding lipid bilayer. This reveals a previously unrecognized mechanism of mechanosensitivity, suggesting that membrane geometry and transporter shape are intrinsically coupled, providing a general principle of regulation across biological systems.

## Introduction

Transmembrane proteins, being embedded in cell membranes, are prone to react to continuous mechanical deformations due to remodeling and stresses applied to the lipid bilayers. Cells use a large family of proteins, the mechanosensitive channels, to control the permeability of their membrane in response to mechanical stresses. This is the case with the eukaryotic ion channels Piezo (1, 2), TRAAK/TREK1 (3, 4), that open at high membrane stretching and allow ions to flow through. It has also been proposed on theoretical basis that these channels could also be sensitive to local high membrane curvature (higher than about 1/50 nm) (5, 6), which was confirmed with the Piezo channel (1, 7). The effect of membrane curvature is expected to be enhanced for proteins with an asymmetrical (conical) shape (5), that locally deform membranes (8, 9). Interestingly, some studies have revealed mechanosensitivity upon stretching of proteins with other identified functions, such as the voltage-gated channel KvAP (10). In fact KvAP has a conical shape (11, 12) and locally bends the bilayer (9, 13), but only the effect of membrane tension on its function has been explored. We expect that other conical transmembrane proteins are potentially mechanosensitive to membrane curvature. Among them, a sub-group of ABC transporters (ATP-Binding Cassette), known as type IV exporters. These transporters have a domain-swapped arrangement with a V-shape, and are known to be flexible trans-membrane proteins that undergo large conformational changes within the bilayer (14–17). Moreover, they could locally deform membranes, according to numerical simulations on the human ABC Pgp (9). These transporters are thus good candidates to test their sensitivity to membrane mechanical properties.

ABC transporters are ubiquitous proteins present in organisms from bacteria to human. They hydrolyze ATP for the transport of a plethora of substrates such as drugs, lipids, ions, peptides (reviewed in (15, 18)). They share a commune topology with two transmembrane domains (TMD), two nucleotide-binding sites (NBD) and two intracytoplasmic domains that join TMDs and NBDs. The structure and function of ABCs have been extensively investigated. Type IV ABCs, such as the bacterial lipid exporter MsbA, the Pgp human multidrug resistance protein, or the non-exporter gated chloride channel CFTR, have a “V-shape” apo-conformation and undergo significant conformational changes during their function. A simplified transport mechanism generally involves a two-state mechanism (Fig. 1A): ATP binding to the NBDs induces their dimerisation and switching of the protein from an “open” state - also called “apo”- (referred to as inward facing state in physiological conditions) to a “closed” state (outward facing state respectively) with a tight contact between the NBDs and where the substrate is released. Upon ATP hydrolysis, the protein switches back to its open state and its V-shape. The extent of the conformational change depends on the ABC transporter, but the gap between NBDs can be as large as a few nanometers (16) (Fig. S1A and S1B). Importantly, two apo-forms with different NBDs separation have been identified on several ABC exporters, such as Pgp (19), MsbA (20) and BmrA (21–23). These states are referred to as “wide-open” and “open” (Fig. 1A) and demonstrate a certain flexibility of these proteins, that may depend on the transporter, its substrate and its lipidic environment. Similarly, different sub-states of the two NBDs have been identified, revealing a sequential ATP hydrolysis (21, 24).

**Figure 1.**
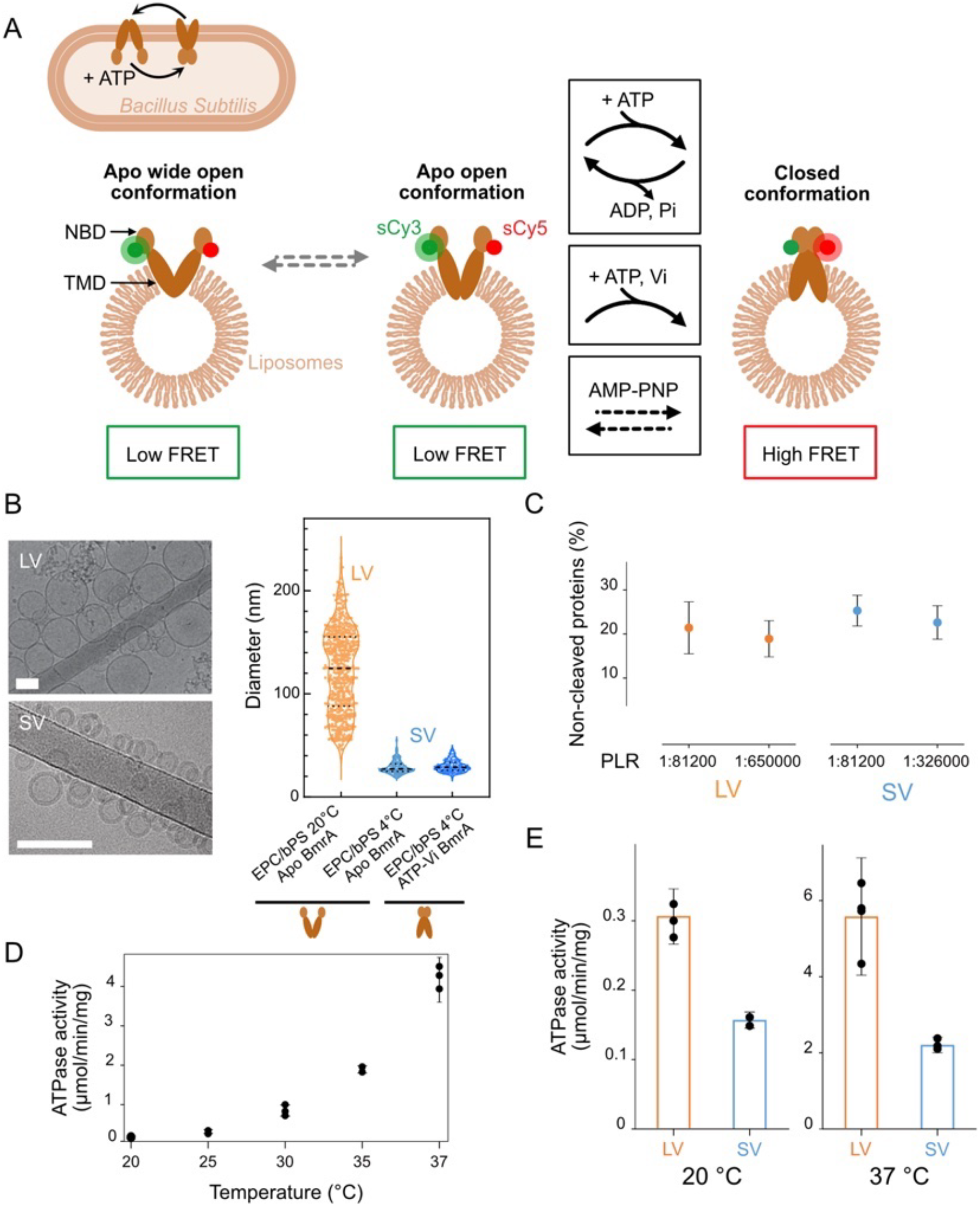
Increasing membrane curvature reduces BmrA ATPase activity. **(A)** Schematic representation of BmrA in *Bacillus Subtilis* (top left) and reconstituted in opposite orientation in liposomes (bottom) in open or apo (inward-facing in bacteria) and closed conformations (outward-facing). The green and red disks indicate the labeling position of sCy3 (donor) and sCy5 (acceptor) for smFRET experiments. The conformational change between apo and closed states occurs actively upon ATP hydrolysis ATP (top box); ATP and Vi covalently trap BmrA in the closed state (middle) while non-hydrolyzable ATP analog, AMP-PNP, binds to NBDs causing reversible protein closing (bottom). **(B)** Left: Cryo-EM images of EPC:bPS (90:10 w/w) apo BmrA proteoliposomes reconstituted at different temperatures: large vesicles (LV) at 20^°^C) and small vesicles (SV) at 4^°^C (scale bar: 50 nm). Right: Size distribution of LV apo BmrA, average diameter ø=123 ± 41 nm (orange, n = 594) and SV with ø=29 ± 6 nm, regardless of BmrA conformation: open (apo (light blue, n=262) or closed (ATP-Vi, dark blue, n=249). **(C)** Protein orientation: Fraction of remaining Alexa 488-labeled BmrA after cleavage of the NBD by trypsin, detected with TIRF microscopy. BmrA was reconstituted in LV and SV at 2 different PLR (number of molecules analyzed from left to right: n = 135, 157, 175 and 168, from 20 movies). **(D)** Effect of temperature on BmrA ATPase activity (in μmol of ATP consumed per min per mg of protein), in detergent (Triton X-100) (3 measurements). **(E)** Effect of liposome size on BmrA ATPase activity measured at 20^°^C and 37^°^C with 5 mM ATP (3 measurements).

In this study, based on its large conformational changes, we have selected BmrA, a homodimeric multi-drug resistance exporter from *B. subtillis*, to study the effect of membrane curvature on its function and conformational dynamics. The structures of BmrA in micelle of detergent are known in the closed form (25), highly similar to the closed conformation of other bacterial exporters, with NBDs in close contact and in the wide-open and narrow-open apo conformations (22). We had previously reconstituted BmrA in lipid membrane at high density and determined a low resolution EM structure, showing an apo conformation with NBDs separated by a few nanometers and a post-hydrolytic state with NBDs in contact (23, 26). This experiment also revealed a major morphological remodeling of the membrane, from curved to planar, between BmrA reconstituted in apo and closed form in high protein density, respectively (26). This demonstrates BmrA’s ability to bend a membrane depending on its state. The apo- and closed-conformations correspond to two different conical shapes of the transporter with a conicity of 15^°^ to 28^°^ (π/12-π/7) in the apo conformation depending on the separation of the NBDs, and an opposite value of -17^°^ (-π/10) in the closed one (Fig. S1C). Altogether, we expect that substantial conformational changes of individual BmrA within the membrane will lead to local membrane deformations, dependent on its conicity during its ATP cycle. Conversely, the conformational changes and function of BmrA should be modulated by membrane curvature.

Practically, snapshots of ABCs’ conformations have been derived with structural techniques (X-Ray, NMR and more recently cryo-EM) (16, 27). Moreover, dynamical conformational changes have been detected on different ABCs, such as the bacterial BmrCD (28), MsbA (19, 29) and McjD (30) or the human Pgp (31–33), CFTR (34) and MRP1 (35) with DEER, LRET or smFRET. Most of these studies were performed on proteins solubilized in detergent micelles or embedded in ∼ 10 nm lipid nanodiscs, where the effect of membrane curvature cannot be addressed. Only four have reported ABC exporters in liposomes, but have not considered the effect of vesicle size (29, 30, 33, 34). Here, we have reconstituted BmrA in Small Unilamellar Vesicles (SUVs) in two populations with small and large diameters. First, we have measured a reduction of ATPase activity for the smallest liposomes. Next, after fluorescent labeling of BmrA on specific NBD residues where the distance is the largest in the apo form, we used TIRF microscopy on immobilized liposomes to detect conformational changes with smFRET. We observed an open (apo) conformation, and a closed one in the presence of ATP or of a non-hydrolysable ATP (AMP-PNP). Interestingly, the fraction of proteins in closed conformation decreases in both cases with increasing membrane curvature, consistent with the observed decrease in activity. Moreover, we propose a theoretical model based on membrane mechanics to quantitatively predict the role of membrane shape and elasticity in the control of the ATPase activity of ABC transporters.

## Results

### BmrA ATPase activity decreases with increasing membrane curvature

We first purified BmrA in detergent (N-dodecyl-β-D-maltoside (DDM)) and reconstituted it in small unilamellar vesicles (SUVs) made of 90% (w/w) Egg Phosphatidyl Choline (EPC) doped with 10% of brain Phosphatidyl Serine (bPS), a negatively charged lipid known to stimulate the ATPase activity of some ABC transporters (36). BmrA (with a molecular weight *M* = 130 kDa for the homodimer) was incorporated at 1:13000 molar (homodimer) Protein Lipid Ratio (PLR). We found that by adjusting the SUVs preparation temperature, we could obtain two different size distributions of the proteoliposomes, which we characterized by cryo-EM (Fig. 1B): i) upon preparation at 20^°^C, a broad distribution of large vesicles “LV”, with a diameter ø = 123 ± 41 nm; ii) upon reconstitution at 4^°^C, a narrow distribution of small vesicles “SV”, with a diameter ø = 29 ± 6 nm. Importantly, the conformation of the reconstituted protein, in apo form or in the presence of ortho-vanadate (Vi) (a commonly used inhibitor that prevents from the NBDs dissociation) had no impact on the size of the small liposomes (ø = 29 ± 6 nm with apo BmrA and ø = 30 ± 6 nm for ATP-Vi BmrA). Moreover, these cryo-electron microscopy images showed that liposomes were unilamellar (Fig. 1B). Finally, we showed that vesicles’ size and unilamellarity remained unchanged after being exposed for a long time to different temperatures (between 20^°^C and 37^°^C) in the course of our experiments described further (Fig. S2).

With floatation in sucrose gradient, we measured 90-95 % protein incorporation in both samples (middle and top fractions) without aggregated proteins in the bottom fraction (Fig. S3). We compared the fluorescence of Alexa 488-BmrA in liposomes, by TIRF microscopy, exposed or not to trypsin that cleaves the NBDs, and we measured that for LV and SV, 80% and 76.15 % of the proteins had their NBDs facing outside the liposomes, respectively, thus accessible upon addition of ATP in the external medium (Fig. 1C). Accordingly, proteoliposomes were similar in lamellarity, amount and orientation of incorporated proteins and differed essentially by their size.

Using a standard enzymatic assay, we measured the ATPase activity *A* of BmrA in detergent micelles and after reconstitution in proteoliposomes in the presence of 5 mM ATP. Note, we performed our experiments without a substrate since BmrA activity is weakly dependent on its presence (18, 37). The activity was 3.8 µmol ATP/min/mg of protein in Triton X-100 at 37^°^C. Surprisingly, we found out that BmrA activity is reduced by a factor 22 when the temperature decreases from 37^°^C to 20^°^C (Fig. 1D), in contrast with previous measurements on Pgp (38) with a factor 4 only on the same temperature range. The activity significantly increased after reconstitution in vesicles, as reported for several ABC transporters (26, 39, 40). Note that, in the following, we have corrected the ATPase activities to account for the fraction of proteins not accessible to ATP, as measured with TIRF microscopy (Table S1). However, after reconstitution (PLR=1:13000 molar), the ATPase activity strongly decreased when reducing the liposome size. Indeed, at 37^°^C, we measured a 2.2-fold lower activity in the highly curved liposomes SV (*A*=2.5 µmol ATP/min/mg of protein), as compared to the larger vesicles LV (*A* = 5.6 µmol ATP/min/mg of protein) (Fig. 1E); this corresponds to a turnover τ = 365 ms in SV and τ = 165 ms in LV for the hydrolysis of two ATP molecules per BmrA molecule, respectively (τ = 2×60.10^3^/*A*×*M*, in seconds, with *M* the homodimer molecular weight). The activity is comparable to previously published data on liposomes containing BmrA (∼6 µmol of ATP/min/mg) (37) or the closely-related ABC transporter MsbA (∼5 µmol of ATP/min/mg) (39). At 20^°^C, a similar curvature-dependence of the activity was also measured, with *A* = 0.31 µmol ATP/min/mg of protein for LV and *A* = 0.15 µmol ATP/min/mg of protein for SV, corresponding to turn-overs τ =2,98 s and τ = 6,15 s, respectively (Fig. 1E). Moreover, by measuring the activity at 37^°^C for ATP concentrations ranging between 0.1 mM to 5 mM, we have confirmed that for both vesicle populations SV and LV, 5mM ATP corresponds to saturating conditions (Fig. S4). Our data show that membrane curvature strongly impacts BmrA activity; however, the protein is able to maintain its cycle in spite of the mechanical stress experienced in the context of the small liposomes. To further decipher the consequences of this strain on the protein conformational changes, we have next investigated the effect of membrane curvature on BmrA structure using single molecule FRET.

### Single-molecule FRET assay to monitor the conformational dynamics of BmrA in LV and SV liposomes

For smFRET experiments, BmrA was labeled with sulfo-Cy3/sulfo-Cy5 (SI Appendix; Fig. S5) on the two native cysteines C436 located at the basis of the NBDs, close to the TMDs, for which the distance change during the protein cycle, thus the FRET change, are expected to be the largest (Fig. 1A, Fig. S1A and S1B). Due to the homodimeric nature of BmrA, 25% of the proteins are doubly labeled (Materials & Methods). The activity of the labeled proteins was conserved at 80% as compared to the non-labeled. Proteoliposomes (containing 0.5% of biotin-PEG-DSPE) were immobilized onto custom-made flow cells, extensively cleaned and pre-treated with silane-PEG-biotin and neutravidin (protocol adapted from (41)). The specificity of the tethering strategy, the integrity preservation of the tethered liposomes and the amount of protein per liposomes (to ensure single molecule regime) were extensively controlled (SI Appendix; Fig. S6).

Our FRET experiments were performed using TIRF microscopy with alternating laser excitation (ALEX) at 532 nm (Donor = sCy3) and 638 nm (Acceptor = sCy5) (42), with an acquisition rate of 1/200 ms^-1^. We measured the intensities *I*_*D*_, *I*_*F*_ and *I*_*A*_ of single molecules over time in the Donor, FRET and Acceptor channels, respectively, and the apparent uncorrected FRET efficiency 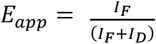 until either the donor or the acceptor bleaches. The sensitivity of our system was checked with a well-established “DNA ruler” assay (43, 44) (SI Appendix; Fig. S7; Table S2). We noted that with the same oxygen scavenger system (Materials and Methods) the dyes’ average fluorescence lifetime was much shorter when linked to BmrA than to DNA, probably due to an enhanced photostabilization of the cyanines dyes upon interaction with the DNA bases (45).

### BmrA exhibits two conformations upon addition of ATP analogs

It is largely reported that type IV ABC exporters switch between two extreme states during the transport cycle: i) a nucleotide-free apo-form where the NBDs are separated by a distance that depends on the ABC and ii) a closed form with dimerized NBDs upon nucleotide binding (e.g. ATP) (14). The lifetime of the closed conformation can be significantly increased by using ATP analogs in the presence of Mg^2+^. Two commonly used ones are AMP-PNP and ortho-vanadate (Vi) (46) (described on Fig. 1A). AMP-PNP is a non-hydrolysable form of ATP, which binds to the NBDs and leads to their dimerization with the protein in a pre-hydrolytic conformation. This binding is reversible, but since it does not involve ATP hydrolysis, the NBDs remain dimerized for a longer time than with ATP. In contrast, in the presence of ATP, Vi is expected to form covalent bonds with the ADP phosphate group bound to the NBD, preventing it from being released and therefore maintaining the protein in a closed post-hydrolytic form and inhibiting its ATPase activity. We first confirmed the effect of both compounds on BmrA in vesicles: BmrA activity in the presence of 5mM ATP is strongly reduced by 80 % upon addition of 2 mM ATP-Vi (well above K_m_=0.05 mM (37)), and by only 25% while adding 5 mM AMP-PNP (Fig. S8A). The inhibition by the ATP-Vi is conserved between 20^°^C and 37^°^C (Fig. S8B).

We have then used our single-molecule assay to measure the FRET efficiency and thus the conformations of single BmrA molecules in the presence of ATP-Vi or AMP-PNP in the SV and LV liposomes, at 20^°^C. In both cases, the analogs were added *prior to reconstitution* at a concentration of 5 mM to the mix of protein/lipid in detergent. In order to reduce dissociation due to nucleotide depletion, the analogs were kept at 5mM in the observation buffer.

In the presence of Vi, we expected to observe mainly a high FRET conformation, corresponding to the closed state of the dimer. Surprisingly, the FRET efficiency histograms exhibited a two-peak distribution in both LV and SV populations (Fig. 2A; 2D histograms with apparent efficiency *E*_*app*_ and stoichiometry *S*_*app*_ are available on Fig. S9; Table 1). This demonstrates the co-existence of two states: i) a high FRET peak at <*E*_*app*_> = 0.45, corresponding to the closed state with an interdye distance of about 4 nm from known structures (25) (Fig. S1); ii) a low FRET <*E*_*app*_> = 0.10, corresponding to an open conformation where the NBDs are separated about 7 nm according to the recent structure (22). Note that, as we suspected, we cannot distinguish between a wide- and a narrow-open conformation, due to the smFRET resolution in this distance range. We thus call this low FRET population “open” (Fig. 1A). From the two-Gaussian fit, we extracted the respective weight of the low and high-FRET populations; we found that they have the same weight for both LV and SV, around 50% (Table 2). Importantly, no transition was detected between the open and closed state on individual time traces (Fig. 2C-D). This result is also enforced by measuring the time average FRET efficiency per protein with the same low and high-FRET populations (Fig. S10). Altogether, this shows that half of the proteins are open, and half are closed during the time of our experiments, independent of the size of the liposomes. Considering the high activity of our BmrA preparation (about 3 times higher than in (47)), this is probably not due to a dead fraction. Moreover, since the open fraction is the same for LV and SV, the effect of membrane mechanics can also be discarded. Note that, although surprising, such a co-existence of open and closed conformation in ATP-Vi has previously been reported, in nanodiscs and detergent, for BmrA in ensemble experiments (48), for its bacterial homolog MsbA (19), human homolog Pgp (32), and for the heterodimer BmrCD (28).

**Table 1.** Two-gaussian fit parameters of BmrA smFRET histograms for the 2 vesicle populations (LV and SV), at 20 ^°^C (RT) et 33^°^C (HT). Amplitude *a*, mean <*E*_*app*_>, and width parameter *w* for the Low-FRET (LF) and High-FRET (HF) peaks

| **RT** |  | **$a_{LF}$** | **$\langle E_{app} \rangle_{LF}$** | **$w_{LF}$** | **$a_{HF}$** | **$\langle E_{app} \rangle_{HF}$** | **$w_{HF}$** |
| --- | --- | --- | --- | --- | --- | --- | --- |
| **LV** | ATP Vi | 0.052 | 0.117 | 0.071 | 0.026 | 0.445 | 0.125 |
|  | AMPPNP | 0.053 | 0.149 | 0.085 | 0.018 | 0.458 | 0.129 |
|  | Apo | 0.077 | 0.119 | 0.087 | 0.005 | 0.40 | 0.139 |
|  | ATP | 0.063 | 0.131 | 0.083 | 0.011 | 0.41 | 0.158 |
| **SV** | ATP Vi | 0.05 | 0.109 | 0.071 | 0.039 | 0.445 | 0.088 |
|  | AMPPNP | 0.054 | 0.15 | 0.094 | 0.014 | 0.4 | 0.141 |
|  | Apo | 0.068 | 0.145 | 0.089 | 0.004 | 0.451 | 0.316 |
|  | ATP | 0.069 | 0.121 | 0.081 | 0.007 | 0.401 | 0.214 |
| **HT** |  | **$a_{LF}$** | **$\langle E_{app} \rangle_{LF}$** | **$w_{LF}$** | **$a_{HF}$** | **$\langle E_{app} \rangle_{HF}$** | **$w_{HF}$** |
| **LV** | ATP Vi |  |  |  |  |  |  |
|  | AMPPNP | 0.045 | 0.095 | 0.07 | 0.023 | 0.400 | 0.154 |
|  | Apo | 0.104 | 0.116 | 0.06 | 0.003 | 0.400 | 0.141 |
|  | ATP | 0.051 | 0.096 | 0.065 | 0.023 | 0.402 | 0.143 |
| **SV** | ATP Vi |  |  |  |  |  |  |
|  | AMPPNP | 0.063 | 0.092 | 0.06 | 0.016 | 0.400 | 0.18 |
|  | Apo | 0.092 | 0.110 | 0.06 | 0.007 | 0.403 | 0.158 |
|  | ATP | 0.068 | 0.079 | 0.06 | 0.016 | 0.400 | 0.16 |

**Table 2.** Fractions of proteins in closed (P_*cl*_) states for the two vesicle populations LV and SV, at 20 ^°^C (RT) et 33^°^C (HT). P_cl_ corresponds to the non-corrected High-FRET fraction calculated from the parameters of Table 1, 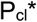 to the fraction corrected of the High-FRET fraction in the corresponding apo state P_cl_(apo).

| RT | Condition | $P_{cl}$ (%) | $P_{cl}^*$ (%) | HT | Condition | $P_{cl}$ (%) | $P_{cl}^*$ (%) |
| --- | --- | --- | --- | --- | --- | --- | --- |
| LV | ATP Vi | $48.4 \pm 0.6$ | $38.6 \pm 4.6$ | LV | ATP Vi | | |
| | AMPPNP | $35.0 \pm 3.9$ | $25.2 \pm 6.0$ | | AMPPNP | $55.4 \pm 1.0$ | $49.1 \pm 2.3$ |
| | Apo | $9.8 \pm 4.6$ | 0 | | Apo | $6.3 \pm 2.1$ | 0 |
| | ATP | $26.3 \pm 1.7$ | $16.5 \pm 4.9$ | | ATP | $52.5 \pm 1.5$ | $46.2 \pm 2.6$ |
| SV | ATP Vi | $51.2 \pm 0.4$ | $35.9 \pm 4.1$ | SV | ATP Vi | | |
| | AMPPNP | $29.5 \pm 3.1$ | $14.2 \pm 5.1$ | | AMPPNP | $45.3 \pm 1.9$ | $28.1 \pm 4.6$ |
| | Apo | $15.3 \pm 4.1$ | 0 | | Apo | $17.2 \pm 4.2$ | 0 |
| | ATP | $22.2 \pm 3.7$ | $6.9 \pm 5.5$ | | ATP | $41.7 \pm 2.2$ | $24.5 \pm 4.7$ |

**Figure 2.**
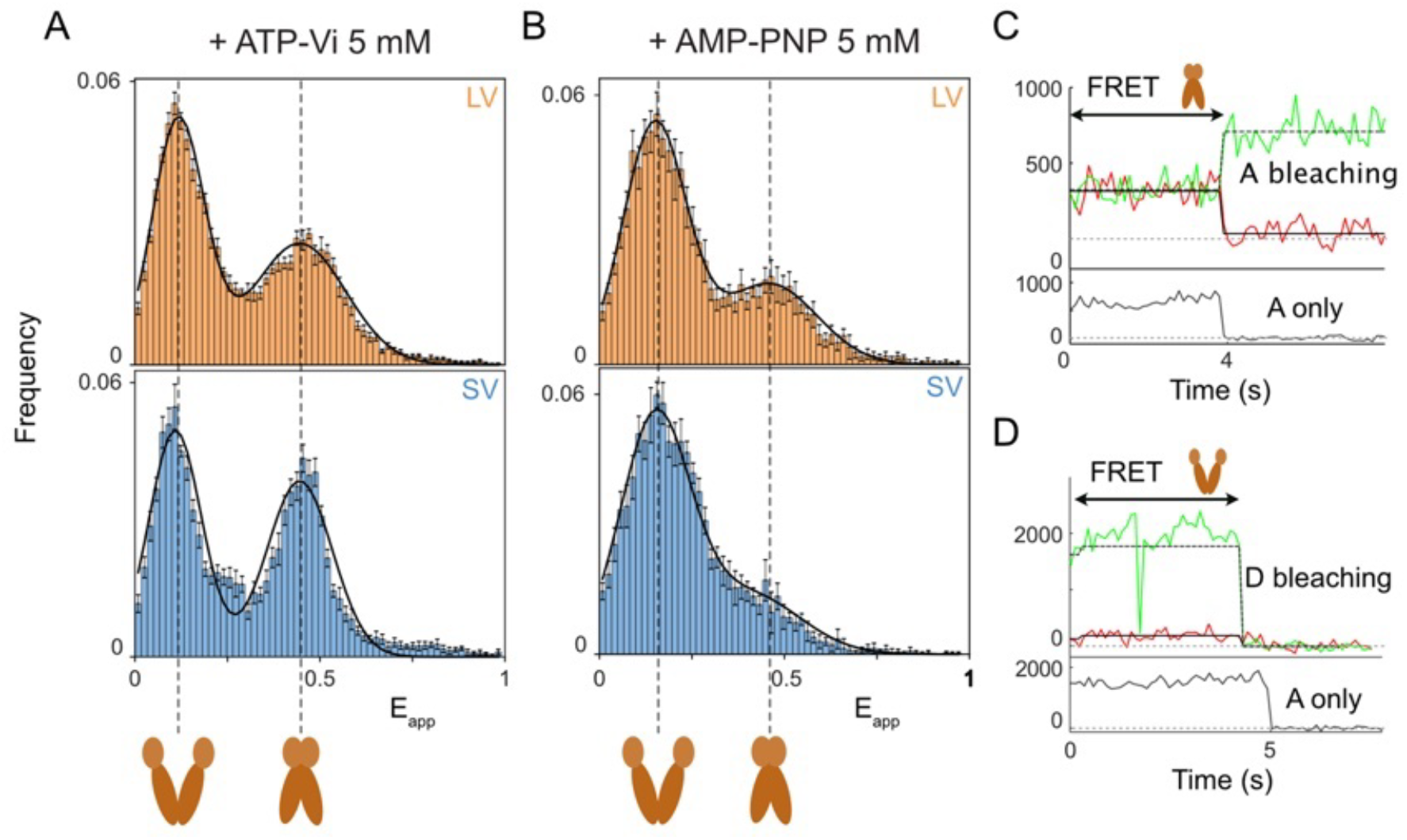
smFRET shows that BmrA closed conformation depends on membrane curvature in the presence of non-hydrolysable ATP. Experiments performed at 20^°^C (RT) **(A**,**B)** Cumulated histograms of apparent FRET efficiency E_app_ upon addition of (A) ATP-Vi 5 mM in LV (top) and SV (bottom) (B) 5 mM AMP-PNP in LV (top) and SV (bottom). The thick line corresponds to the two-gaussian fit. *N*_*all*_: number of time points in histograms, *N*_*prot*_: corresponding number of proteins. (A top) *N*_*all*_ = 18559 and *N*_*prot*_ = 525. (A bottom) *N*_*all*_ = 7552 and *N*_*prot*_ = 325. (B top) *N*_*all*_ = 5109 and *N*_*prot*_ = 157. (B bottom) *N*_*all*_ = 5094 and *N*_*prot*_ = 141. For each condition, data come from two independent experiments. Error bars are calculated using bootstrapping. **(C**,**D)** Typical time traces of proteins in LV: (C) with ATP-Vi 5mM in closed state and (D) in open state. Top panel: donor (green) and FRET (red) intensities over time. Plain and dashed line represent a fit of the FRET switches. Bottom panel: intensity of acceptor (A) only (black).

For proteins reconstituted in the presence of 5 mM AMP-PNP, we also observed a bimodal distribution with peaks at similar positions, showing the co-existence of the same open and closed states, with no switch between these conformations (Fig. 2B; Table 1; Fig. S9 and S10). However, the relative fractions of these states are different from the ATP-Vi condition. First, the fraction in the closed state *P*_*cl*_ is lower in AMP-PNP (35% in LV and 29.5% in SV) (Table 2) than in presence of ATP-Vi (about 50%), in line with our activity measurements in detergent showing a stronger inhibition by ATP-Vi (Fig. S8). Similar differences between these analogs were reported on human ABC Pgp (32, 49) or ABCB7 (50). Second, in contrast with ATP-Vi, the fraction in a closed state is reduced in SV as compared to LV. Altogether, this suggests that, in the presence of AMP-PNP where closed and open forms are in equilibrium, the NBDs dimerization rate depends on membrane curvature, which is not the case when NBDs irreversibly dimerize in complex with ATP-Vi.

### The fraction of BmrA in closed conformation decreases with membrane curvature

We then investigated the conformational changes of BmrA during its ATPase cycle. Considering the strong dependance of the activity with temperature, we performed the smFRET on SV and LV at 20^°^C (RT) and at 33^°^C (HT) (the highest possible temperature for these experiments). We first assessed the apo-BmrA conformation (no ATP) in LV and SV (Figs. 3A-B, top histograms; Fig. S9; Table 2). As expected, we observe a peak at low FRET for both vesicle sizes. Although these distributions would be fitted with a single Gaussian (Fig. S11), a better fit was obtained (notably for SV) with a double Gaussian while fixing the position of a second peak at 0.45, as measured above with the ATP analogs. In this case, we found that a large majority of proteins (90.2 % and 93.7% for the LV at RT and HT, 84.7% and 82.8% for the SV at RT and HT, respectively (Table 2)) are in the open conformation. At both RT and HT, we thus observed a slightly more pronounced high FRET tail for the SV than for the LV, which might correspond to a higher fraction of the inside-facing proteins in SV (Fig. 1C), also more constrained in a closed conformation by the liposome geometry. The low FRET peak is similar to the state observed under addition of ATP analogs (Table 1). This demonstrates that apo-BmrA exhibits an open conformation in vesicles, independent of membrane curvature.

**Figure 3.**
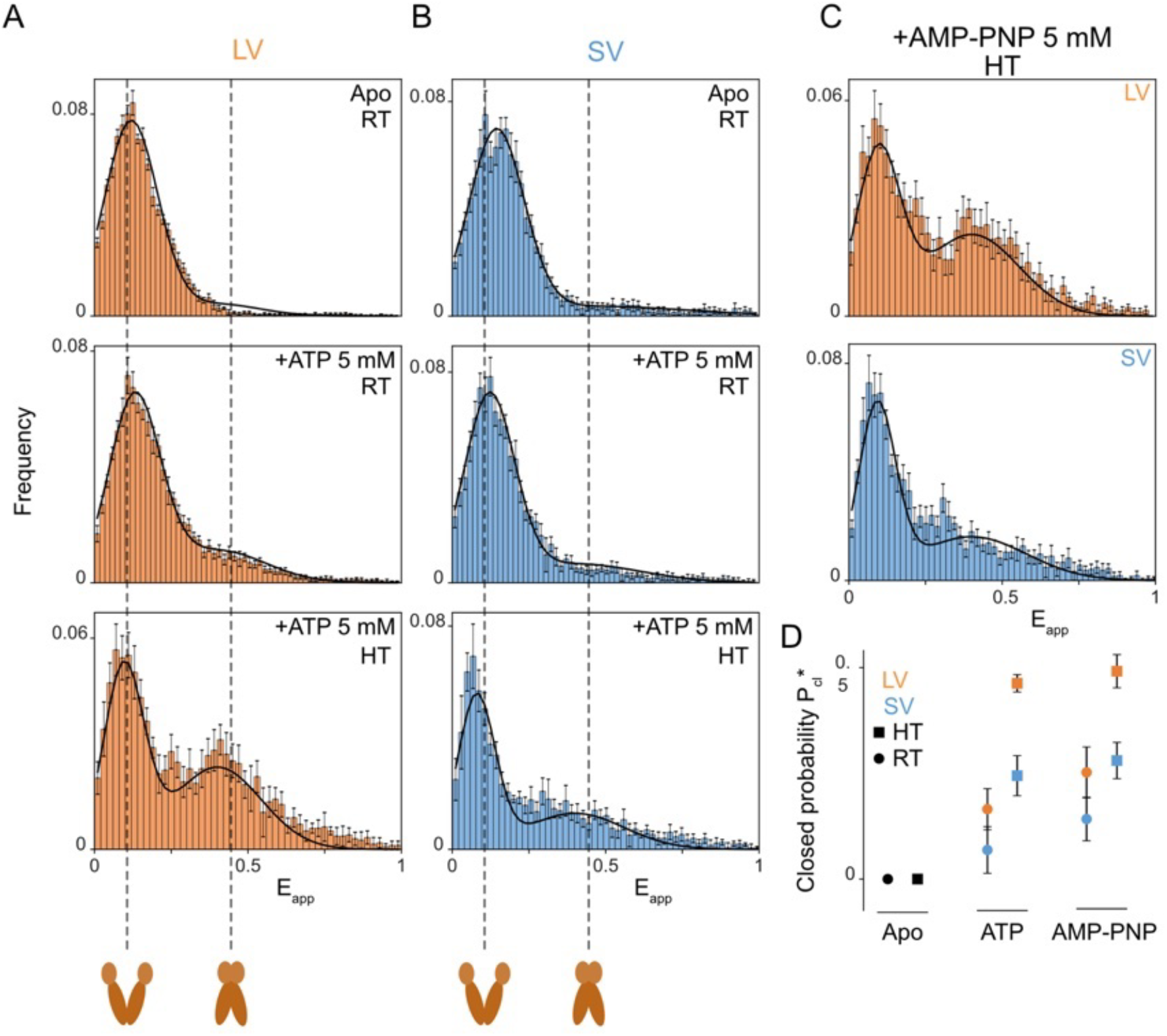
Effect of curvature on conformation of BmrA. **(A**,**B)** Cumulated histogram of *E*_*app*_ for BmrA in apo conformation in LV (top A) and SV (top B) at RT (20^°^C), with 5 mM Mg-ATP in LV (middle A) and SV (middle B) at RT and with 5 mM Mg-ATP in LV (bottom A) and SV (bottom B) at HT (33^°^C). The thick lines show a two-Gaussian fit with the high-FRET peak set to <*E*_*app*_>_HF_ in ATP-Vi. Top A: *N*_*all*_ = 10451, *N*_*prot*_ = 310. Middle A: *N*_*all*_ = 7743, *N*_*prot*_ = 226. Bottom A: *N*_*all*_ =2991, *N*_*prot*_ =1050. Top B: *N*_*all*_ = 2797, *N*_*prot*_ = 115. Middle B: *N*_*all*_ = 3959, *N*_*prot*_ = 130. Bottom B: *N*_*all*_ = 3235, *N*_*prot*_ =1093. Data come from two independent experiments and three for SV apo. Error bars are calculated using bootstrapping. **(C)** Cumulated histogram of *E*_*app*_ for BmrA in LV (top) and SV (bottom) with 5 mM AMPPNP at HT. (top) *N*_*all*_ = 2759, *N*_*prot*_ = 823. (bottom) *N*_*all*_ = 2935, *N*_*prot*_ = 980 **(D)** Probability for proteins to be in closed state 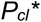 as a function of nucleotide, temperature and liposome size.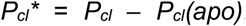, with *P*_*cl*_*(apo)* the high FRET fraction detected in the corresponding apo state. Error bars from bootstrapping errors on *P*_*cl*_ (Table 2).

We finally studied the effect of ATP on the conformational distribution. Interestingly, upon addition of 5mM Mg-ATP, we observed differences between the two vesicle populations, at both temperatures (Figs. 3A-B, middle and bottom histograms; Fig. S12). As above, we analyzed the FRET distributions with a double Gaussian and by setting the high-FRET peak at 0.45 (Table 1). The position of the peak corresponding to the open form remains essentially unchanged. However, as compared to the apo condition, we detected a second high-FRET population corresponding to the closed form, with a fraction P_cl_ that depends on the SUV size and on the temperature. Addition of ATP resulted in an increase of closed protein population of 46.3% at HT and 16.5 % at RT, as compared to the apo form for the LV (Fig. 3A, middle-bottom histograms; Table 2). But the variation of the fraction of closed proteins in SV was only 24.5% at HT and 6.9% at RT (Fig. 3B, middle-bottom histograms; Table 2), significantly lower than for the LV. Consistent with our activity measurements, higher at HT than at RT, the relative fraction of closed BmrA is higher at HT, for both SV and LV (Figs. 3A-B, bottom histogram; Table 2). Similarly, with the AMP-PNP analog, we also observed an increase of the relative probability for the protein to be in closed conformation when experiments are performed at HT (Fig. 3C). In conclusion, we show here that, both at RT and HT, high curvature reduces the probability of the protein to be in a closed state in the presence of ATP or of AMP-PNP (Fig. 3D).

Despite the oxygen scavenger present in our buffer, the FRET traces lasted 7 seconds on average before bleaching at 20^°^C (Fig. S13A and S13B) and only about 1.5 seconds at 33^°^C due to an increased phototoxicity with temperature (Fig. S14). Though a rare observation, we detected a switch between both conformations on some traces at 20^°^C (Fig. S13C and S13D), where ATP-induced conformational dynamics is visible). This is consistent with the very slow turn-over measured with our ATP-ase activity assay (3 s for LV and 6.2 for SV). Such switches were not observed at 33^°^C.The analysis of the traces (Fig. S13A and S13B) and the time-averaged FRET histograms (Fig. S15) at 20^°^C show that in the presence of ATP, BmrA proteins are either open or closed for a few seconds. By measuring the average lifetime of the proteins in the high-FRET state in LV (with *E*_*avg*_ > 0.3, n = 38 proteins /226), we can conclude that in these conditions and at 20^°^C, the lifetime of the closed state is longer than 7 seconds of the same order as the average turnover measured in cuvette. Similarly, at room temperature, lifetime of a few seconds was found for the closed state of McjD (30) and more recently of the human ABC CFTR (34).

### Modeling the effect of membrane curvature on the open to close transition probability

We have developed a model to quantify the influence of membrane mechanics on the cycle of conformational change of BmrA. We assume that the change of protein conformation can be described by a single reaction coordinate, the V shape angle *θ* (Fig. 4A). In the absence of nucleotide, the protein energy landscape *E*_*p*_(*θ*) is assumed to have a minimum in the open conformation *θ*_*op*_, which we model for simplicity as a quadratic energy with a stiffness *A*: *E*_*p*_(*θ*) = *A*(*θ* − *θ*_*op*_)^2^ (Fig. 4B). Thermal fluctuations can bring the protein to its closed conformation *θ*_*cl*_, and if nucleotides are bound to the two NBDs, the landscape gains a deep minimum at *θ*_*cl*_ characterized by a binding energy *ε* when the NBDs bind to one another. We may consider two types of nucleotides, ATP or the non-hydrolysable AMP-PNP. With ATP, the binding energy is assumed to be so large that the protein remains in the closed conformation until ATP is hydrolyzed, which occurs at a prescribed rate *k*_*ATP*_. With AMP-PNP, the system is considered to be at thermodynamic equilibrium, and the closed-open transition occurs when thermal fluctuations bring the protein over the energy barrier *ε*. Membrane mechanics modify the transition rates through the energy of membrane deformation *E*_*m*_(*θ*) associated to the protein shape changes. This term depends on the membrane’s bending rigidity and on the geometry of the liposome in which the protein is embedded (Fig. S16). Membrane shape optimization can be performed analytically for small deviations from a spherical shape (details in SI Appendix). In this case, 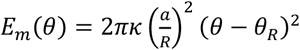, where *κ* is the membrane’s bending rigidity, *a* and *R* are the protein and SUV radii, and the optimal opening angle is *θ*_*R*_ = *sin*^−1^(*a*/*R*) ≈ *a*/*R* if the SUV is in the optimal state and have a spontaneous curvature 2/*R*. In the thermal equilibrium case, the probability ratio of open *P*_*op*_ and closed conformation *P*_*cl*_ follows a Boltzmann statistic. In the non-equilibrium case (when ATP is hydrolyzed), this ratio has a steady-state value given by the balance between the thermal flux (from the open to the closed conformation) and the active reversed flux. Writing Δ*E*_*i*_ = *E*_*i*_(*θ*_*cl*_) − *E*_*i*_(*θ*_*op*_), we find:

**Figure 4.**
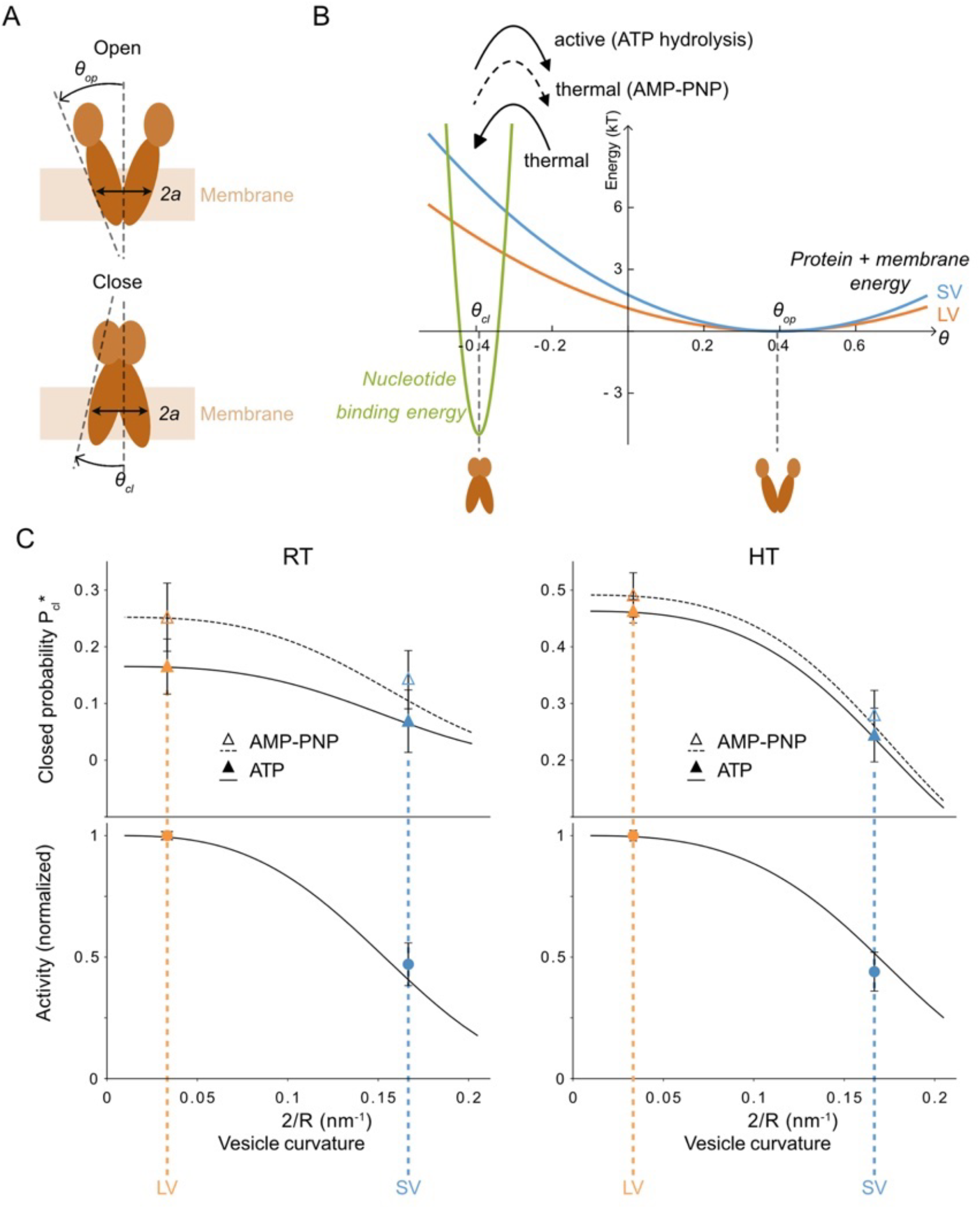
Theoretical 2-states model describing the effect of membrane mechanics on protein shape transition. **(A)** Definition of geometrical parameters *2a, θ*_*op*_ and *θ*_*cl*_ used in the theoretical model. **(B)** Two-state model scheme: energy landscape for BmrA alone favors the open state. With nucleotides, NBDs dimerize, and adding energy (green) favors the closed state. Membrane energy adds a contribution that further disfavors the closed state, more so for SV (blue) than LV (orange). Open-to-closed transition is thermal, while closed-to-open is either thermal (AMP-PNP)) or ATP hydrolysis-triggered. **(C)** Model results: probability of being in open conformation (top, with ATP or AMP-PNP) and activity (bottom, ATP turnover rate), as a function of SUV curvature, at RT and at HT. Experimental data (dots) from 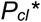 using smFRET (top) and from ATPase assay (bottom). Both data fitted with a single parameter *K≈(27*.*5±6*.*5)kT*. Error bars: noise at high FRET value in Apo (top, Table 2) and standard deviation from measurements (bottom).

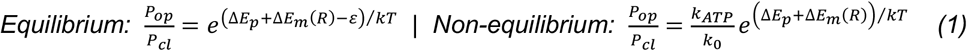

where *kT* is the thermal energy and *k*_0_ is a typical rate of thermal fluctuation of the protein conformation. In the non-equilibrium case, one can also derive the rate of ATP hydrolysis per protein 1/*τ* as:

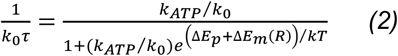

Through the energy difference

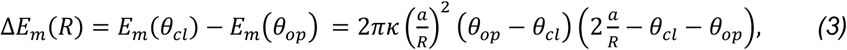

Eqs.(1,2) provide the size-dependence of the open and closed populations, as well as the rate of ATP hydrolysis. As the open conformation is facing outward, we expect *θ*_*op*_ > 0 and *θ*_*cl*_ < 0 and Δ*E*_*m*_ > 0 for liposomes with large curvature (Fig. S1C). Therefore, we expect the closed conformation to be disfavored, and the rate of ATP hydrolysis to be lower in SV than in LV. Note that a proper description of the experimental observation must account for the distribution of vesicle size (SI Appendix; Fig. S17), which we found not to affect significantly our conclusions.

One important prediction of the model is that the open-to-close probability ratio for the two distributions of vesicle size are related by a Boltzmann factor that only involves the membrane deformation energy 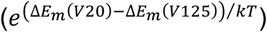 both in the equilibrium (AMP-PNP) and non-equilibrium (ATP) case. This prediction agrees with our observations (Fig. 4C, top; Fig. S17A), with Δ*E*_*m*_(*SV*) − Δ*E*_*m*_(*LV*) ≃ (0.84 ± 0.2)*kT*. Based on BmrA and homologs’ structures, we used a protein diameter 2*a* ≈ 4*nm* (Fig. S1D). Since the conicity of the closed conformation is independent of the protein membrane environment (19, 32), *θ*_*cl*_ was set to −*π*/10 measured from closed BmrA structure in detergent (Fig. S1C). Conversely, it was reported that the opening of the apo form of homologs MsbA and Pgp changes between detergent and nanodisc environments (19, 32). Therefore, we used *θ*_*op*_ ≈ + *π*/10, corresponding to MsbA, structure in membrane (nanodisc), in good agreement with the angle measured for BmrA in vesicles at high protein density but low resolution (23) (Fig.S1A and S1C). We have fitted all our data at both temperatures with these quantities (Fig. 4C, top), using the closed fractions 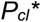 from Table 2. The membrane bending rigidity is the only fitting parameter and is found to be *κ* ≈ (27.5 ± 6.5)*kT* which is in line with expected values (51). Note that Δ*E*_*m*_/*kT* is very small for large SUVs (≃ 6 · 10^−3^) so that proteins are not sensitive to membrane elasticity, while the open-to-closed transition should be moderately impaired in the smaller SUVs (Δ*E*_*m*_ ≃ 0.84*kT*), as observed.

The predicted value of the bending rigidity *κ* strongly depends on the chosen values of *θ*_*op*_ and *θ*_*cl*_. The fit gives access to the value of the energy difference *ΔE*_*m*_ (Eq.3), which increases with *θ*_*op*_ if *θ*_*op*_ is larger than the optimal opening angle fixed by the vesicle size *θ*_*R*_, as is the case for the full range of liposome size we explored. Therefore, the predicted value of the bending rigidity decreases if we use a larger value of *θ*_*op*_. Using the open angle *θ*_*op*_ ≈ + *π*/7, deduced from the BmrA structure in detergent (Fig. S1C), and under the assumption of symmetric states (*θ*_*cl*_ = −*θ*_*op*_) for which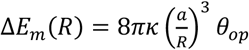, we find a membrane bending rigidity *κ* ≈ (19 ± 4)*kT*, when fitting to the population and activity curves.

Knowing the membrane parameters from the previous fit, we can determine the value of the relevant combination of protein parameters in the active case for the chosen ATP concentration: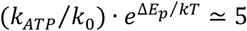. This lets us predict, without any adjustable parameter, how the protein activity (the rate of ATP hydrolysis) depends on the liposome size (Fig. 4C, bottom; Fig. S17B), at RT and at HT. In the case of our experiments, we predict that it is reduced by about 50% for the small SUVs as compared to the larger ones, at RT and HT (Fig. 4C, bottom), a value in very good agreement with the experimental values obtained from ensemble measurements (Fig. 1E).

## Discussion

In general, mechanosensitive membrane proteins have been identified through the cell response after mechanical stresses and then, the corresponding molecular mechanism linked to their function studied at the structural level. Previous studies on MsbA (29) and on ABC importers OpuA (52) and ButCD (53) have highlighted the effect of the local environment for ABC transporters, showing at the single-molecule level different degrees of opening of the apo conformation when incorporated in detergent or in lipid nanodiscs. Our smFRET study reveals that membrane curvature also impacts the conformation changes of an ABC transporter, as well as its activity. Our experiments, in which we control the curvature of liposomes, demonstrate that the activity of BmrA decreases when increasing membrane curvature, as well as the fraction of proteins in a closed conformation. We have also observed that BmrA activity decreases strongly between 37^°^C - close to the optimal growth temperature of *B. subtillis* - and 20^°^C, much more than expected based on Arrhenius law. The effect of curvature, both on the ATPase activity and on the conformation changes, at two different temperatures, can be recapitulated with a single theoretical model based on the idea that the protein conformation changes are associated to membrane deformation and are therefore sensitive to membrane mechanics.

Regarding the open conformation, a wide range of openings have been reported among ABC exporters, with NBDs in close contact up to NBD separated by about 6-7 nm (14, 16, 27, 54). Our smFRET data in vesicles are consistent with the recent 3D model of BmrA in detergent (22) with dissociated NBDs in the apo form or MsbA in E. coli bacteria (20). However, we have not detected any difference for the low FRET between LV and SV, in contrast with the transition rate between both conformations. Former studies on bacterial MsbA and human PgP stressed that this relative opening is wider in detergent than in lipid nanodisc (19, 32), suggesting that the protein environment influences the structure of these transporters.

Although the differences between SV and LV are significant, a more precise test of our model would require data points between 60 nm down to 20 nm where the effect of curvature is expected to be observed for BmrA; experiments where the single molecule FRET traces would be measured simultaneously with the corresponding liposome diameter deduced from membrane fluorescence (55) would be optimal, though challenging in single molecule experiments due to fluorescence leaking.

As in our in vitro experiments with BmrA NBDs pointing outward, the curvature of the liposome membrane fits the open conformation better than the closed one, membrane energy then favors the open state. In cell, the orientation of BmrA is inward facing with the NBDs toward the cytoplasm, so that the effect of membrane energy should favor the transition from open to closed conformation; thus, we could expect an increase of the protein’s activity during the formation of extracellular vesicles from the plasma membrane or the formation of vesicles in the lumen of compartments (ILVs), but a reduction in intracellular vesicles that have an opposite topology.

Our model predicts that such an effect of membrane curvature can be expected for all type-IV ABC exporters, provided that the apo form is sufficiently open, since they have similar size and well-conserved closed conformations. From the model, we infer that ABCs with BmrA characteristics are mechanosensitive only at large membrane curvature, higher than 0.08 nm^-1^, i.e. for vesicles smaller than 50 nm diameter and for tubules, 25 nm diameter. In *Bacillus subtilis*, such high curvature is reported upon the formation of extracellular vesicles (56), where the activity of the transporter could be enhanced due to the physiological orientation of the protein and the geometry of the vesicles. The human Pgp being a close homolog to BmrA (Fig. S1), we expect that membrane curvature does not impair its function in usual transport vesicles inside cells (typically 50 to 60 nm diameter), endosomal compartments or the Golgi (57), where it has been reported to be present in addition to the plasma membrane. Thus, although potentially mechanosensitive, the structural design of this transporter might ensure its proper functionality under physiological conditions. Nevertheless, significant effects should be detected at weaker curvature for proteins with a larger membrane imprint (*a*) and/or with higher conicity (*θ*_*op*_ and *θ*_*cl*_) (Fig. S18). Membrane bending rigidity (κ) can also modulate the conformational switch (Eqs. (1-3)). Consequently, changing membrane stiffness by adjusting lipid composition (e.g. via cholesterol or sphingomyelin concentration) is expected to tune the sensitivity of the activity to membrane curvature; however, lipid composition by itself can also affect protein activity (58).

Altogether our results demonstrate that membrane curvature plays a role in the transport cycle kinetics of ABC transporters, which should depend on protein lateral size, conicity and membrane stiffness. This invites us to re-think the definition of *mechanosensitivity* to also include other proteins whose activity involves large shape changes within the membrane. BmrA’s sensitivity to membrane bending suggests a potential response to membrane tension. However, curvature and membrane tension could have different consequences. While the effect of membrane tension on mechanosensitive proteins has been largely documented, it would be worth to investigate the effect of tension on the function of this new class of mechanosensors.

## Materials and Methods

### Reagents

Egg Phosphatidylcholine (EPC), brain Phosphatidylserine (bPS), 1,2-distearoyl-sn-glycero-3-phosphoethanolamine-N-[biotinyl(polyethylene glycol)-2000] (DSPE-PEG(2000)-Biotin) and 1,2-dihexadecanoyl-sn-glycero-3-phosphoethanolamine, triethylammonium salt (Texas Red® DHPE) were purchased in CHCl3 stock solution from Avanti Polar Lipids (Alabaster, AL, USA); BODIPY®-FL-C5-HPC from Invitrogen; N-dodecyl-β-D-maltoside (DDM) and Anapoe® X-100 (Triton® X-100) from Anatrace (Maumee, OH, USA). 50 mesh BioBeads® were purchased from Bio-Rad (Hercules, CA, USA) and prepared according to (59). Lacey carbon electron microscopy grids were from Ted Pella (Reding, CA, USA). Nucleotides, ATP, AMP-PNP, ATPase inhibitor ortho-vanadate, pyranine, neutravidin, biotin-BSA (from bovine milk > 98%), D-glucose, glucose oxidase, Trolox and all other chemicals were purchased from Sigma Aldrich (Merck KGaA, Darmstadt, Germany); SulfoCy3 and SulfoCy5 maleimide probes from Lumiprobe GmbH (Germany); Coverslips (thickness # 1) from Menzel-Gläser (Germany); Silane-PEG2000 and silane-PEG3400-biotin (in powder) from LaysanBio (Alabama, USA) and further dissolved in DMSO; double stranded DNA for FRET calibration from Eurofins Genomics (Ebersberg, Germany); Catalase from bovine liver from MP Biomedicals (California, USA).

### BmrA expression, purification and labeling

His-C-ter-BmrA was expressed in *E. coli* and purified in 0.05 % DDM as previously reported(23). The elution buffer from Ni-NTA agarose beads contained 50 mM Tris (pH 8.0), 100 mM NaCl, 250 mM imidazole (pH 8.0), 10% glycerol and 0.05% DDM. Proteins were eluted at 0.3-1.5 mg/mL (Bradford assay) without further concentration. Purified proteins were aliquoted in 50 µL volume, stored at -80^°^C, and thawed only once before use.

For smFRET experiments, purified BmrA from the elution step was fluorescently labeled with maleimide dyes on C436, the sole Cys residue of each monomer (two per homodimer). BmrA in detergent was incubated overnight at 4^°^C (pH 7) with sulfo-Cy3 and sulfo-Cy5 maleimide dyes (selected as the most suitable, see SI Appendix), at a molar ratio of sCy3/sCy5/protein=10/10/1 (Fig. S5). Unbound dyes were removed by rebinding BmrA at pH 8 on Ni-NTA agarose beads in batch, followed by washing with 5 volumes of a dye-free buffer and elution with 250 mM imidazole. Labeled BmrA was analyzed on SDS-PAGE gel by fluorescence and Coomassie blue. Labeling efficiency was determined from the concentration of the dyes, measured by absorbance of sCy3 (molar extinction coefficient *ε*_*max*_ = 162 000 cm^-1^.M^-1^), sCy5 (*ε*_*max*_ = 281 000 cm^-1^.M^-1^), and from the protein concentration (Bradford assay). Labeling yields were 32% for sCy3 and 39% for sCy5. BmrA being a homodimer, we expect that statistically, the labeling probabilities are: unlabeled 8%; single-labeled sCy3 19%; doubly-labeled sCy3/sCy3 10%; single-labeled sCy5 23%; doubly-labeled sCy5/sCy5 15%; doubly-labeled sCy3/sCy5 25%. ATPase activity was measured as described below, and sCy3/sCy5 labeled BmrA retained 80% of the unlabeled protein.

### Reconstitution of BmrA in proteoliposomes

Proteoliposomes containing BmrA were formed by detergent removal using Bio-Beads® from a fully solubilized lipid/protein/detergent mix, as described previously (37) and according to the principle reviewed in (60, 61). Pure lipid liposomes composed of EPC:bPS (90:10 w:w) and 0.5% of biotin-PEG2000-DSPE were prepared by drying a lipid film, resuspending at 10 mg/mL in water, and sonicating with a tip sonicator for 1 min on ice. The suspension was aliquoted (50 µL) and stored at –20 ^°^C.

For reconstitution, one aliquot was diluted to 2 mg/mL in R buffer (50 mM MOPS, 150 mM NaCl, 1 mM MgCl_2_, pH 7.5), and solubilized with Anapoe® X-100 for 15 min at a detergent/lipid ratio of 2.5 (w:w). Note that BrmA is stable and functionally active both in Anapoe® X-100 and in DDM. After lipid solubilization, for smFRE, BmrA was added at protein-to-lipid ratios (PLR) of 1:650000 for LV and 1:325000 for SV (SI Appendix). For AMP-PNP and ATP-Vi experiments, inhibition was induced by adding either 2 mM Vi + 5 mM ATP, or of 5 mM AMP-PNP, both in the presence of 5 mM MgCl_2_. After 15 minutes, detergent was removed by adding BioBeads® under gentle stirring, at 20^°^C (LV) and 4^°^C (SV). BioBeads® were added in three successive rounds: i) for LV (20^°^C), 10 w:w (2h), 10 w:w (1h) and 20 w:w (1h); ii) for SV(4^°^C), 30 w:w (2h), 30 w/w (1 h), 60 w/w (1 h), the higher amounts ensuring comparable detergent removal. Proteoliposome suspensions were collected and stored at 4 ^°^C for up to 24 h. During this period ATPase activity remained stable, and vesicle morphology was preserved as confirmed by cryo-EM.

### Protein incorporation and orientation in proteoliposomes

Protein incorporation was measured after separation of proteoliposomes, aggregated proteins and protein-free liposomes by sedimentation in a sucrose gradient according to a protocol modified for proteoliposomes(62). BmrA was reconstituted at PLR=1:9750 in EPC-PS 9/1 mol/mol and 2 mg/mL lipid concentration at 20^°^C or 4^°^C. Floatation assay was performed with polycarbonate centrifugation 11x34 mm tubes (Beckman Coulter, USA) and sucrose solutions prepared in HK buffer (50 mM Hepes, pH 7.5, 150 mM NaCl). An aliquot of 150 µL of proteoliposomes was gently mixed with 100 µL of 60 % sucrose solution. Then, 200 µL of a second layer of 24% sucrose was added followed by 100 µL of HK buffer. Tubes were ultracentrifuged 90 minutes in a TLS-55 Swing bucket rotor (Beckman) at 55,000 rpm at 4^°^C with a slow acceleration and deceleration steps (5/5). Fractions were gently pipetted off with a syringe and analyzed by 4-12% NuPage gel after staining with Commassie Blue. Protein intensities and incorporation yield were measured with the Amersham Imager 680 and the analysis Software, version 2.0.0.

Protein orientation in the membrane was determined from single molecule fluorescence measurements. Alexa-488 labeled BmrA was reconstituted in large and small proteoliposomes at PLRs of 1:81200 and 1:326000 for SV, and 1:81200 and 1:650000 for LV. For immobilization, samples were prepared as described below. The fraction of proteins with NBDs pointing outward was measured by comparing the number of immobilized fluorescent molecules in standard samples versus samples incubated with Trypsin (0.1 mg/mL) for 30 minutes. Molecule detection was performed using the MATLAB code iSMS(63, 64), described below.

### ATPase activity

ATP hydrolysis measurement was carried out using a coupled enzyme assay of pyruvate kinase, and lactate dehydrogenase by absorbance changes of NADH/NAD+ ratio at 340 nm(37). All measurements were made in triplicates. ATPase activity was measured in 50 mM MOPS pH 7.5, 150 mM NaCl, 10 mM of MgCl_2_ with 1X enzyme cocktail and 0.5-1 μg of protein at 37^°^C in 150 μL total volume. Absorbance of NADH was measured by spectrometer (Cary Win 60 UV – Vis, Agilent Technologies) at 340 nm every 15 s with quartz cuvette (N.105-254-15-40, Hellma Analytics, 3 x 3 mm). Proteoliposomes were incubated for 4 min at desired temperature. ATP was added at 1.5 or 5 mM final concentration and absorbance was recorded for 4 min (first minute of reaction was ignored for ATPase activity calculation). Since the *K*_*D*_ of ATP for BmrA is 1 mM, this ensured that BmrA hydrolyzed ATP at maximum rate in all conditions(37). Rate of inhibition was measured after addition of 2 mM of orthovanadate or 5 mM of AMP-PNP. ATPase activity was calculated by quantity of NADH oxidized equivalent to quantity of ATP hydrolyzed, per min and per mg of protein. ATP hydrolysis of BmrA in micelles of detergent was measured with Anapoe® X-100 0.05% in the cuvette. The amount of BmrA per assay was 1 μg for proteoliposomes and 2 μg for BrmA in detergent.

The activity at 33 ^°^C was inferred from the measurements of activity in both detergent and vesicles. First the activity in 0.05% detergent was interpolated from the measurements made at 30^°^C and 35^°^C (Fig. 1D). Then, using an average proportionality factor between LV activity and detergent activity of 1.65 (1.8 at 20^°^C and 1.5 at 37^°^C), we estimated the activity in LV at 33^°^C (2.48 µmol ATP/min/mg, corresponding to a turnover τ = 329 ms). Finally, using again the average proportionality factor between SV and LV activity at both 20 ^°^C (2.1) and 37 ^°^C (2.5), we calculated the activity in SV at 33^°^C (1.1 µmol ATP/min/mg, corresponding to a turnover τ = 764 ms).

### Cryo-electron microscopy and size measurement of proteoliposomes

Holey formvar lacey grids (Ted Pella, USA) were glow discharged for 30 s under vacuum. Then, 4 μL of 1 mg/mL of proteoliposomes were deposited during 30 s on the grid before flash frozen in liquid ethane with an EM-GP plunger (Leica Microsystems, Germany). Images were taken under low dose condition on a Glacios electron microscope (Thermofisher, USA) operating at 200 KV and recorded with a Falcon IVi camera. Diameter of proteoliposomes were measured manually from cryo-EM images from 594, 262 and 249 vesicles reconstituted at 20^°^C, 4^°^C with Apo BmrA and 4^°^C with ATP-Vi BmrA.

### Proteoliposomes immobilization for smFRET experiments

A small channel of 10 µL was custom-built between two coverslips (Menzel-Gläser, thickness # 1): bottom coverslip is 25 mm diameter, and top squared-shaped coverslip is cut to 18×10 mm^2^. The coverslips were cleaned by sonication with the following sequence: chloroform (20 min, then dried under the hood for 5 min), water (5 min), 1M KOH (20 min), three rinsed with water, and a final wash in water (15 min). They were dried under a gentle nitrogen stream and exposed to air plasma for 2 min. The observation chamber was assembled using a double-sided tape mask. To specifically tether proteoliposomes, the chamber was incubated with a silane-PEG solution containing 1% of silane-PEG-biotin (5 mM, 1hr); rinsed with 100 µL of R buffer; neutravidin (40 µL at 20 µg/mL, 5-10min); rinsed with 100 µL R buffer; β-casein (40 µL at 1 mg/mL,10 min); rinsed, and finally, proteoliposomes (40 µL, 5.10^-2^ mg/mL and 5.10^-3^ mg/mL lipids for LV and SV, respectively, 5–10 min) before a last rinse with 100 µL R buffer and imaging.

### Microscopy and smFRET assay

Observation was performed on a Nikon TI-Eclipse microscope equipped with a Nikon TIRF arm, a TIRF oil objective ×100 (NA 1.45, Nikon), a DualView system (Photometrics DV2) and an ORCA-Flash4.0 V3 Digital CMOS camera (Hamamatsu). The DualView system was equipped with a dichroic mirror 633 nm (ZT633rdc, Chroma) and the appropriate filters for the FRET couple sCy3/sCy5 (593/40 and 684/24 respectively, Chroma). We used alternating laser excitation (ALEX) at 532 nm (sCy3) and 638 nm (sCy5 (Lasers Oxxius). Acquisition rate was 1/200 ms^-1^, with an exposure time of 50 ms in each channel and a 50 ms time-lag in between (necessary to overcome delay issues in our system).

To heat the smFRET samples at 33^°^C, we inserted the chambers into a heating unit Tempcontrol 37-2 Digital (PeCon GmbH, Germany). To limit evaporation, excess of buffer was kept on the inlet and outlet of the chamber.

All experiments were carried out utilizing an oxygen scavenging system supplemented with 1 mM Trolox as a photostabilization agent(41). The oxygen scavenger system consists of 4 mg/mL D-glucose, 1 mg/mL glucose oxidase and 0.04 mg/mL catalase; it was prepared at the last minute before observation and filtered at 0.2 µm before to load it in the observation chamber (SI Appendix). Note that the oxygen scavenger system was more effective on the DNA sample, allowing us to observe minutes-long intensity traces, compared to the single-BmrA observation, for which the average fluorescence lifetime was 7 sec (SI Appendix). Attempt to improve the photostability of the markers with another oxygen scavenger (PCD/PCA) was unsuccessful.

### smFRET analysis

Data analysis was performed with the Matlab code iSMS(63, 64), for detecting positions of Donor and Acceptor, matching pairs Donor/Acceptor and to measure intensities I_D_, I_F_ and I_A_ respectively in the Donor, FRET and Acceptor channels, according to time (all intensities are corrected for background). The automatic bleach detection was used but could be manually adjusted if necessary and blinking intervals were set manually. In the software, the data evaluation process was then performed manually on Donor/Acceptor pairs. We discarded events with more than one bleaching step in either channel, traces that showed very rapid photobleaching, or interference of strongly fluorescent neighboring peaks (resulting in highly noisy intensity trace) were also excluded at this point. For each single molecule, the apparent FRET efficiency *E*_*app*_ *= (I*_*F*_*) / (I*_*D*_ *+ I*_*F*_) and the apparent stoichiometry *S*_*app*_ *= (I*_*D*_ *+ I*_*F*_*) / (I*_*D*_ *+ I*_*F*_ *+ I*_*A*_) were measured until either the donor or the acceptor bleaches, and were accumulated into a normalized histogram, without applying further corrections. All histograms were plotted using a number of bins set to the average of √*N*, where N is the number of points in each individual histogram. Gaussian fits were performed using the *minimize* function from the Python module *scipy*.*optimize*, with the constraint that the fit had to be normalized, in accordance with the histogram. The following equation is used for the Gaussian fit on apparent FRET efficiency 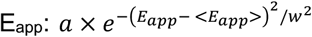 with *a* the amplitude, *<E*_*app*_ *>* the mean position and *w* the parameter related to the Gaussian width. When double Gaussian fit was applied, the parameters of the fit were used to measure the probability to find the protein in low FRET state (= *P*_*op*_) or in high FRET state (= *P*_*cl*_). Errors were measured by applying bootstrapping on our data (10 times).

### Theory

See theoretical section of the SI Appendix.

## Acknowledgments

We thank Elie Balloul and Nikos Hatzakis (University of Copenhagen) for support for the data analysis and insightful discussions. We thank Julien Maufront for analysis of the cryo-images of proteoliposomes. We thank Bassam Hajj, Julien Pernier, Xavier Solinas (Ecole Polytechnique), Simon L’Horset (Centrale Supélec) for technical support and David Lacoste (ESPCI), Peter Tieleman, Estefania Barreta Ojeda (University of Calgary) for providing conceptual ideas.

This work was supported by the Agence Nationale pour la Recherche (ANR-16-CE11-0026-01 for DL, MD and PB; ANR-19-CE11-0002-01 for RKS and PS). AD was funded by the Sorbonne University, the Doctoral school “Physique en Ile de France” (ED-564), the Institut Curie and the grant ANR-16-CE11-0026-01839. KB is funded by the “École normale supérieure Paris-Saclay” doctoral program. SJP was funded by Doctoral school ED “Complexité du vivant” (ED-515) and the University Paris Sciences et Lettres. PB team is supported by the Fondation pour la Recherche Médicale (FRM) (FRM EQU202003010307). PB is member of the CNRS consortium AQV. AD, SJP, MD, PS, DL and PB are members of the Labex Cell(n)Scale (ANR-11-LABX0038) and Paris Sciences et Lettres (ANR-10-IDEX-0001-02). We acknowledge the Cell and Tissue Imaging Core facility (PICT IBiSA), Institut Curie, member of the French National Research Infrastructure France-BioImaging (ANR10-INBS-04).

## Supporting Information for

### Supporting Information Text

#### Single-molecule FRET - Additional details and control experiments

##### Labeling

In order to fluorescently label the homodimeric BmrA for smFRET experiments (Fig. 1A), we used the two native cysteines C436 located at the basis of the NBDs, close to the TMDs, for which the distance change during the protein cycle, thus the FRET change, are expected to be the largest (Fig. S1A and S1B). Several couples of donor/acceptor dyes were evaluated according to the rate of labelling and the ATPase activity of the labelled proteins including Alexa488/Atto610, Alexa488/sulfoCy5, Cy3B/sulfoCy5 and sulfoCy3/sulfoCy5 selected based on spectrum overlap, dye photostability and published data. Best results have been obtained with sulfoCy3/sulfoCy5 in terms of photochemistry, labelling ratio, and protein activity preservation (Fig. S5). The activity of the labelled proteins was conserved at 80% as compared to the non-labelled protein.

##### Reconstitution

During the reconstitution at 4^°^C for the SV preparation, the amount of BioBeads® was increased to ensure that the rate of detergent removal was similar than at room temperature. These amounts of BioBeads® were sufficient to remove all detergent molecules including the amount of DDM present in the preparation of BmrA (1). As also shown previously, at this low PLR, negligible amounts of lipid and no protein were adsorbed onto the BioBeads® (2).

##### Oxygen scavenger preparation

A 10x D-glucose stock solution can be prepared and stored at -20^°^C for several months. A 100x “gloxy” solution (mixture of catalase and glucose oxydase) was prepared and stored at 4^°^C for up to several months. The Trolox 2x stock solution was prepared at approximately 1-2 mM, with pH adjusted to 7.5, filtered at 0.2 µm and stored in the dark at 4^°^C up to two weeks. Solutions were mixed to a final 1x concentration just before before observation (solution was filtered a last time at 0.2 µm before to load it in the observation chamber).

##### smFRET

Our FRET experiments were performed using TIRF microscopy with alternating laser excitation (ALEX) at 532 nm (Donor = sCy3) and 638 nm (Acceptor = sCy5) (3). We measured the intensities *I*_*D*_, *I*_*F*_ and *I*_*A*_ of single molecules versus time in the Donor, FRET and Acceptor channels, respectively, with an acquisition rate of 1/200 ms^-1^. Upon following selection criteria (described in Materials & Methods), we retained data points corresponding to molecules labelled with both a donor and an acceptor, for which we measured over time the apparent uncorrected FRET efficiency 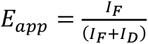 until either the donor or the acceptor bleaches (an oxygen scavenger solution was used to limit bleaching). We then plotted the distribution of instantaneous *E*_*app*_ values from multiple traces (total number of proteins N_prot_ > 100), and performed a single or double Gaussian fit, extracting a peak mean position *<E*_*app*_*>* and a width parameter *w* (equation and parameters are defined in Material & Methods). The sensitivity of our system was checked with a well-established assay (4, 5) (See below; Fig. S7; Table S2), using three 40 base-pair double-stranded (ds) DNAs, carrying the same donor/acceptor pair, at distances ranging between 3 and 7 nm, similar to interdye distances on BmrA and its homologs (Fig. S1A and S1B).

Single molecule experiments require to have maximum one protein (homodimer) per liposomes. In practice, we aim for having about one tenth of liposomes containing a protein, reducing the probability to have two proteins per liposome to 1%. To determine the concentration of protein required to be in this regime, we have incorporated sCy5-labelled BmrA in fluorescently labelled SV liposomes with 0.01% of BODIPY®-FL-C5-HPC lipid, which allowed detecting simultaneously the liposomes and the proteins (Fig. 6A). For PLR= 1:81200, we found that 40% of vesicles co-localized with sCy5-BmrA while 60% did not contain any protein. For PLR= 1:406000, only 5% of the liposomes contained a protein. We have thus reconstituted the transporter at PLR= 1:326000 in the SV, and PLR= 1:650000 in the LV.

In our FRET assay, the liposomes containing 0.5% of biotin-PEG-DSPE, were immobilized onto custom-made flow cells, extensively cleaned to get rid of fluorescent contamination (Fig. S6B) and pre-treated with silane-PEG-biotin and neutravidin (Fig. S6C). The specificity of this tethering strategy was confirmed since very few liposomes are bound to the surface in the absence of neutravidin (Fig. S6D). Typical microscopy images of Donor and FRET channels are depicted on Fig. S6E. We established that the bound vesicles remained sealed and non-permeable to small molecules by encapsulating the soluble dye pyranine in the liposomes labelled with 0.5% of Texas Red® DHPE (Fig. 6F). Co-localization of both fluorescent signals indicated that up to 80% of liposomes were intact after tethering to the surface. We found that the percentage of intact liposomes decreased when increasing the PEG-Biotin density on the surface, due to membrane ripping open. We selected a PEG-Biotin density of 1% for our experiments.

##### smFRET assay calibration using DNA rulers

The system was thoroughly tested with a 40 base-pair double-stranded (ds) DNA as a ruler (Fig. S7A) (4, 5). One strand was labelled with a biotin molecule on the 3’ end, allowing to immobilize the DNA on a coverslip, pre-treated with BSA-biotin and neutravidin (see below), and on the 5’ end with a Cy3 dye. The complementary strand was modified subsequently on three different T-bases with a Cy5 dye, resulting in three distances *L*_*D-A*_ between the donor and acceptor fluorophores, respectively 19, 12 and 7 base-pairs, corresponding to 6.7, 4.7 and 3.4 nm (distances calculated using the Inter-Dye Distances Helical Model of DNA (4)).

The oligonucleotides are hybridized at a final concentration 15 µM in each oligo, in buffer 10 mM Tris-HCl pH 8, 400 mM NaCl, 1 mM EDTA (buffer TE). The solution is brought to 84.6^°^C for 10 min, then allowed to cool down to RT. Glass coverslips were extensively cleaned and then built together as described in Material & Methods. A solution of biotinylated BSA at 1 mg/mL is incubated in the chamber for 5 min and rinsed with 100 µL of buffer TE. The surface is coated with neutravidin at 20 µg/mL for 5 min, rinsed with 100 µL of buffer TE, incubated with β-casein at 1 mg/mL for 10 min and rinsed with 100 µL of buffer TE. Just before observation, the DNA sample is loaded at 30 pM for 2 min and rinsed with 100 µL of buffer TE.

Single peak histograms were measured for each sample (Fig. S7B) and a single Gaussian fit was performed to obtain the mean *<E*_*app*_*>* value (respectively the width parameter *w*), namely 0.23 (resp. 0.12), 0.51 (resp. 0.13) and 0.87 (resp. 0.12) for the 19, 12 and 7 bp constructs, respectively (see Table 1 for the parameters of the fits; Fig. S7C; example traces displayed on Fig. S7D and S6E). This calibration proves that the sensitivity of our system is suitable for detecting BmrA conformational changes, since for BmrA homologs, the distance between our labeling position varies between about 4 and 9 nm, depending on the species and the protein environment (Fig. S1 and S7C).

##### Mathematical description of our model. (separate file)

Modeling the effect of membrane curvature on the open to close transition probability

**Fig. S1.**
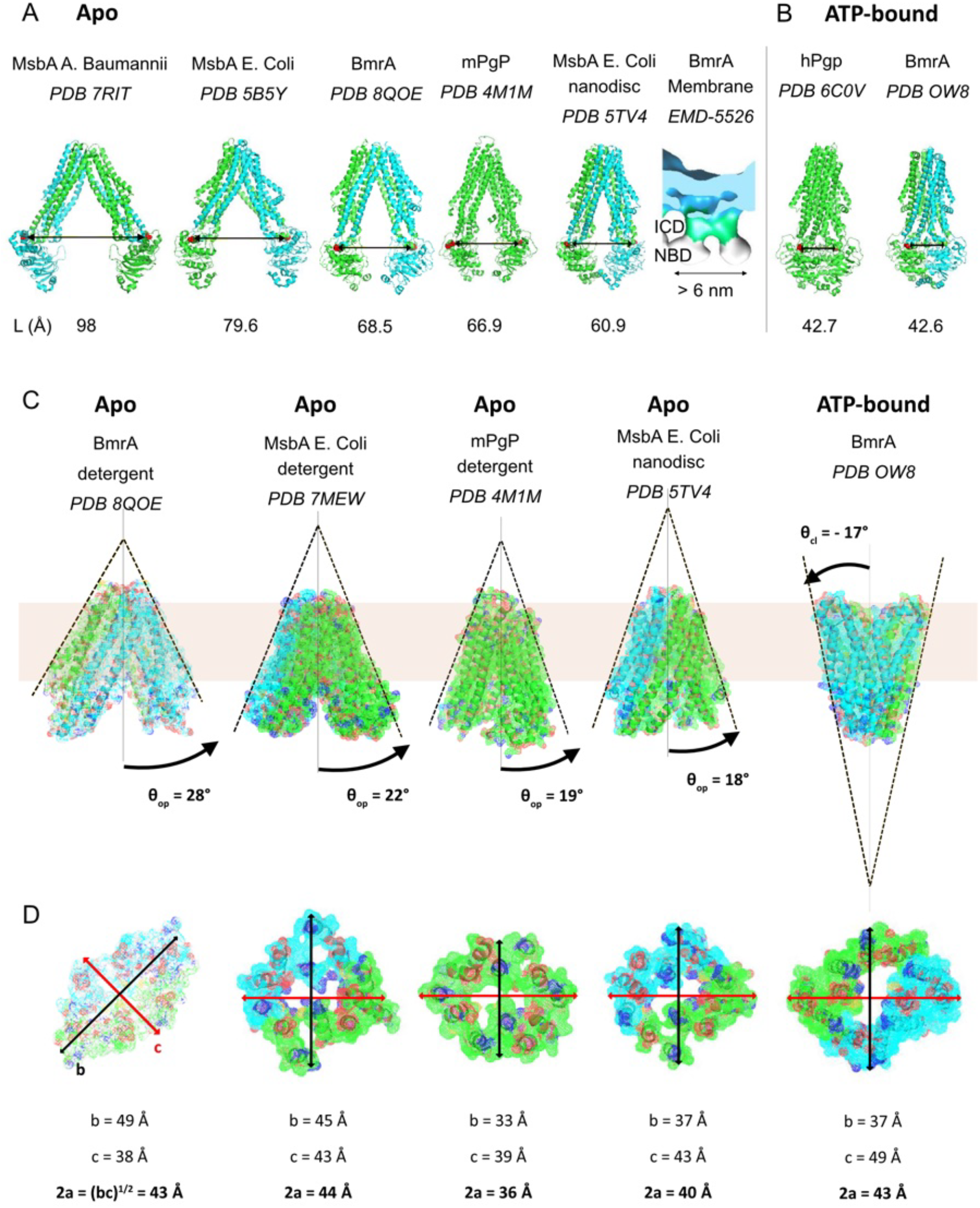
ABC exporters exhibit a large range of opening in the apo form but similar NBD organization in closed conformation. Structures from RCSB PDB (https://www.rcsb.org/) and EMDB (https://www.ebi.ac.uk/emdb/) of BmrA and ABC exporters sharing high homology with BmrA: bacterial MsbA and human and mouse Pgp. **(A)** In apo state. Distance L (in Å) between red-labelled residues, equivalent to those used for BmrA labelling (C436) and PDB ID. For BmrA, the 3D cryo-EM model in detergent and the 23 Å resolution cryo-electron microscopy density map in vesicles (NBDs: white, intracytoplasmic domain ICD: green, TMD and lipid bilayer: blue) are depicted. **(B)** In closed state in the presence of ATP for human Pgp and BmrA. **(C)** Assessing the protein geometrical parameters for the model. Side view of ABC transporters transmembrane domains. The V-shape angle of the open form θ_op_ is shown for apo BmrA in detergent and for ABC exporters sharing high homology with BmrA: MsbA and Pgp. Similar angles for Pgp in detergent, MsbA in detergent and MsbA in nanodiscs. The V-shape angle of the closed conformation θ_cl_ is measured on the structure of ATP-bound BmrA. Membrane is represented in beige. For simplicity, NBDs have been removed from the 3D models. **(D)** Top view of the transmembrane domains at the membrane middle plane. The protein diameter 2a is deduced from 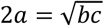 where b and c are the ellipse radii, in the middle plane of the membrane.

**Fig. S2.**
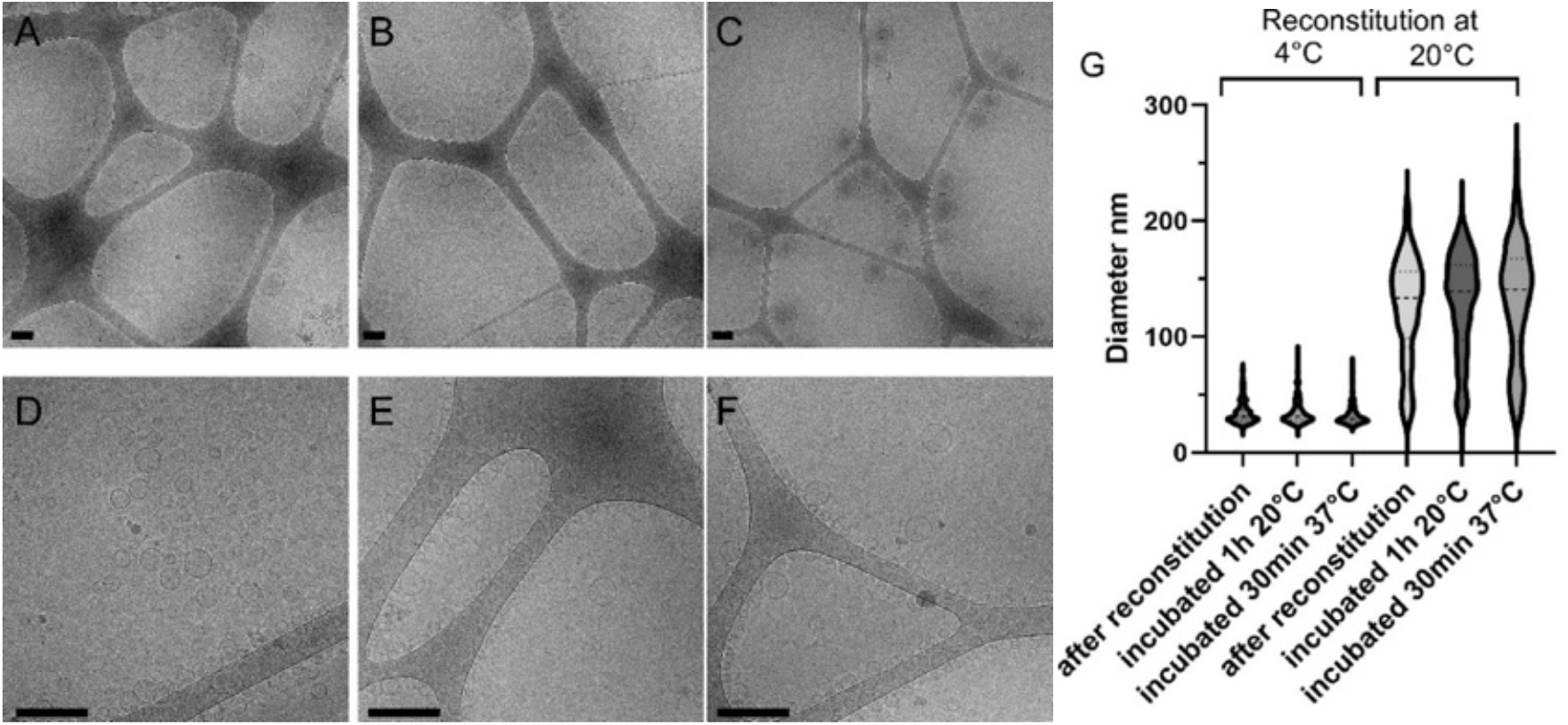
Cryo-EM images of proteoliposomes reconstituted at 20^°^C (A, B, C) and at 4^°^C (D, E, F). Samples have been flash-frozen 20 min after reconstitution (A, D), after 1h additional incubation at 20^°^C (B, E), and after 30 min additional incubation at 37^°^C (C, F). Bars: 50 nm. (G) Violin graph of vesicles’ diameters from A (n=226), B (n=215), C (n=301), D (n=277), E (n=254), F (n=318).

**Fig. S3.**
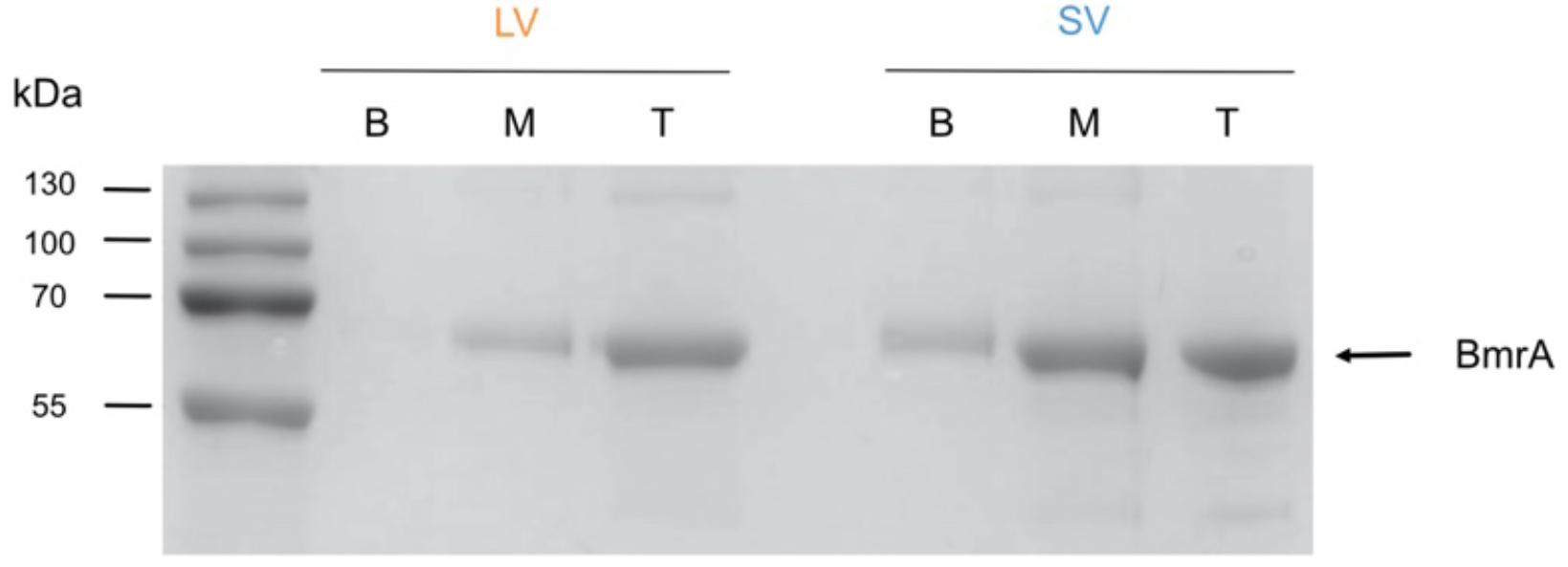
Floatation assay in a sucrose gradient. It first shows BmrA incorporation in LV (left) and SV (right). Bottom (B), middle (M), and top (T) fractions were analyzed on a gel, indicating minimal protein aggregation and proper BmrA incorporation.

**Fig. S4:**
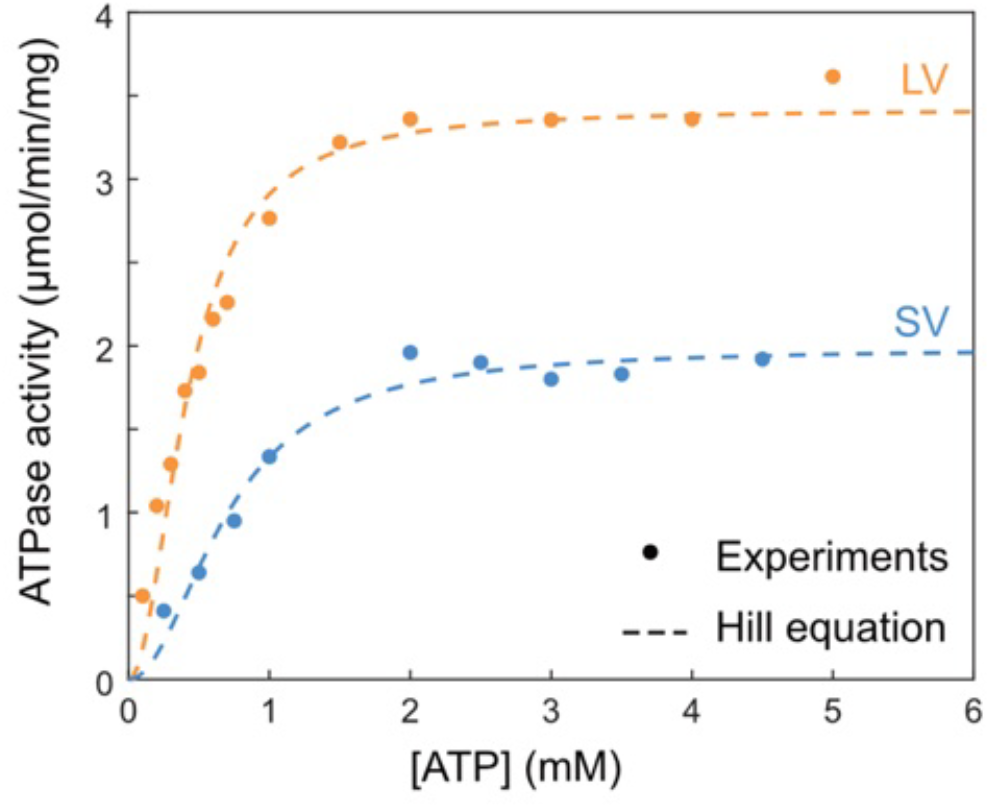
Measure of the ATPase activity as a function of ATP concentration for SV (blue) and LV (orange). K_m_ and V_max_ are deduced from the fit to the Hill equation with a coefficient 2, since 2 ATP molecules are involved in the process: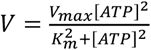. SV: K_m_=0.7 mM, V_max_ =2,0 μmoleATP/mg/min; LV: K_m_=0.4 mM, V_max_=3.4 µmole ATP/mg/min. This shows that 5mM ATP corresponds to saturating conditions for both LV and SV. Using a smaller value for the Hill coefficient does not change our conclusion about the saturating conditions. Moreover, K_m_ is lower for LV than for SV, in qualitative agreement with our model where the transition rate between open and close state depends on the liposome size.

**Fig. S5.**
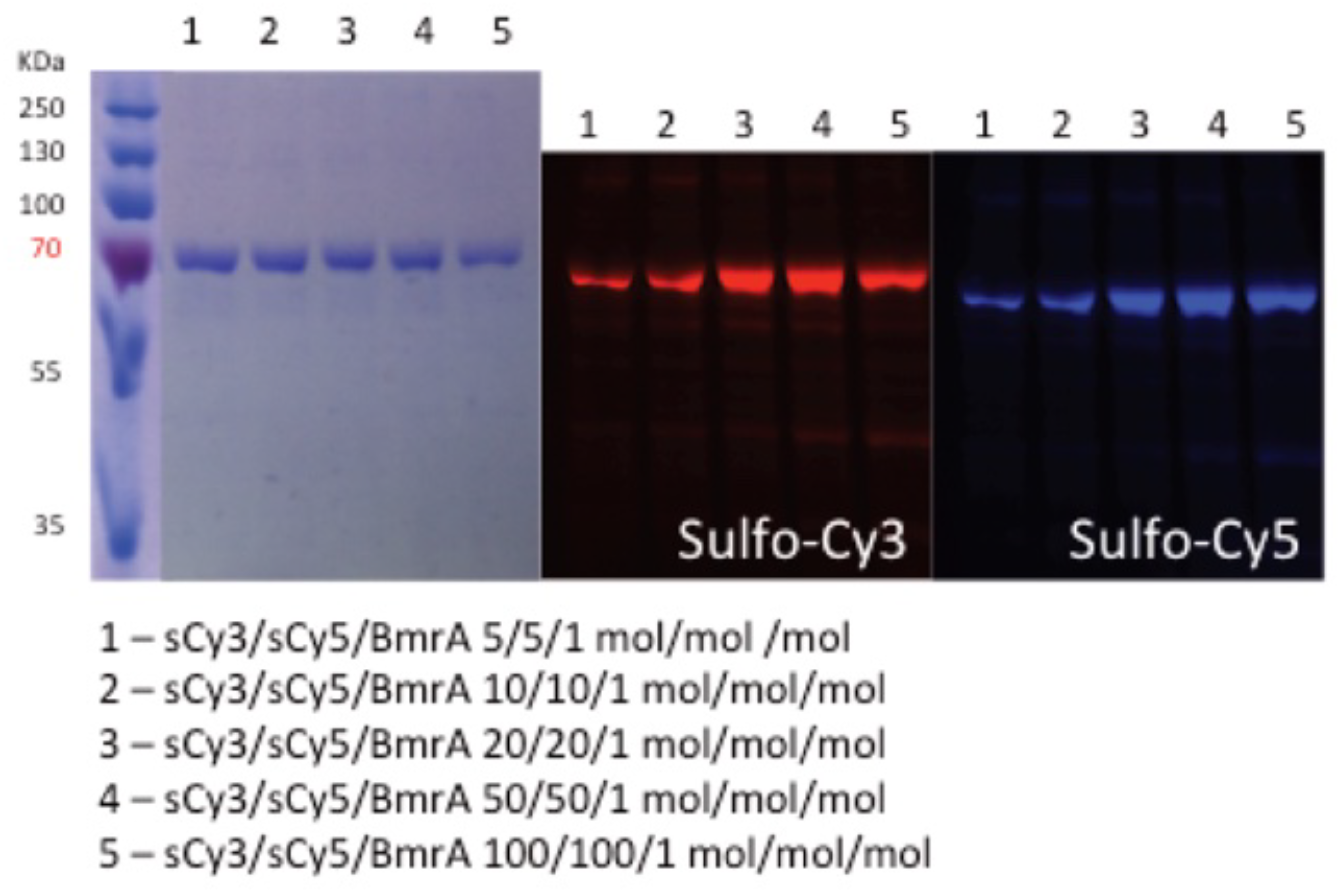
Protein labelling optimization. NuPage gel of BmrA. solubilized in DDM. labelled with increasing amount of sCy3 and sCy5 dyes for optimizing the protein double-labelling, at 4^°^C overnight incubation. Condition 2 was selected as a compromise between labeling efficiency and activity preservation.

**Fig. S6.**
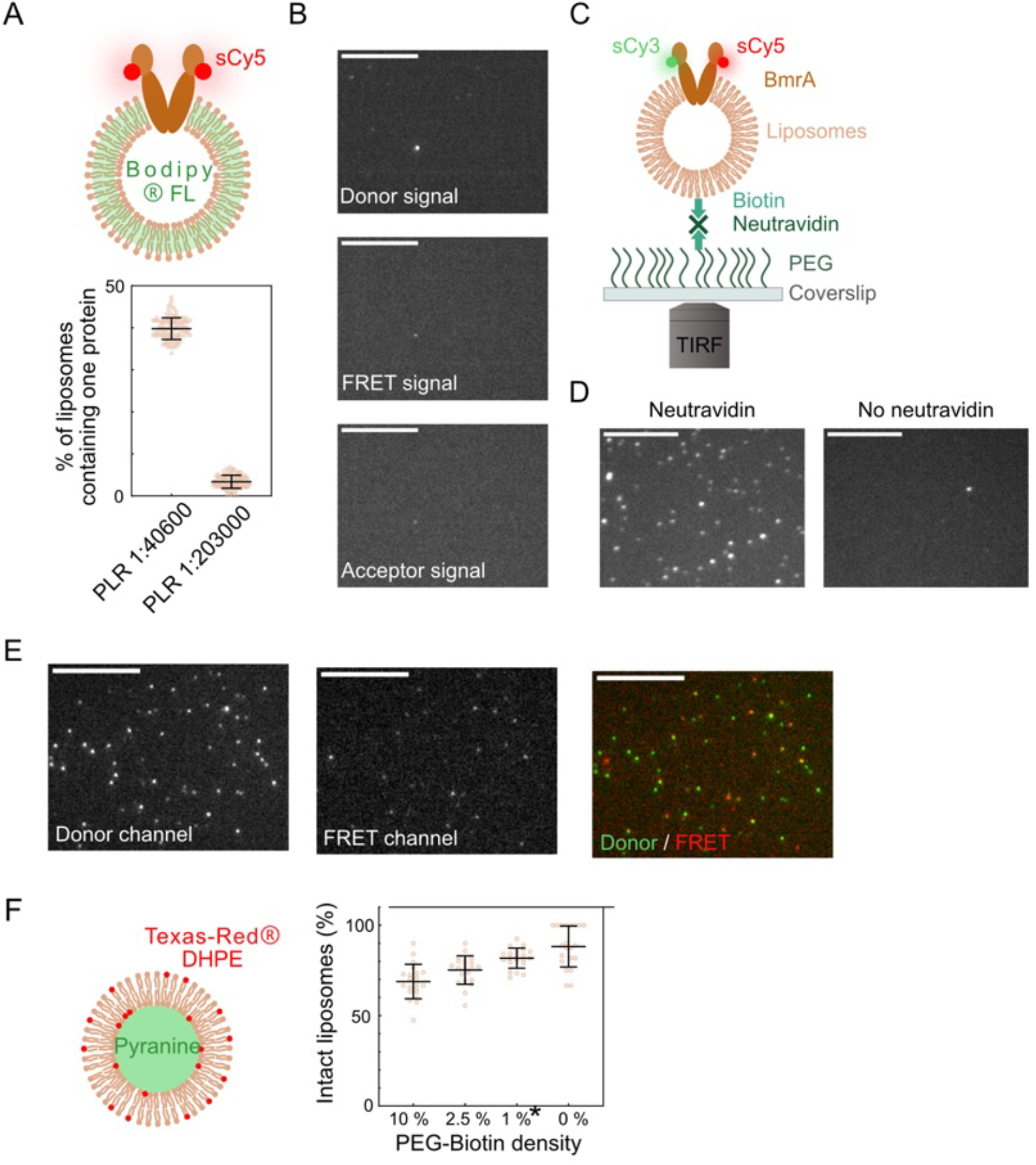
Control experiments for smFRET. **(A)**Control of the single molecule regime in SV, using a double labeling of the protein and of the liposome membrane. Top: Scheme of the experiment. sCy5-labelled BmrA is reconstituted in SV that contain 0.01% fluorescent lipids (Bodipy®-FL). Colocalization of sCy5 and Bodipy®-FL signals indicates a liposome containing at least one protein. Bottom: Percentage of liposomes (per image) containing at least one BmrA, for two different PLR, 1:203000 and 1:40600. In both cases, most liposomes are empty. 120 images were analyzed. **(B)**Control for the cleanliness of surfaces. Surface observation after the cleaning process and surface treatment (full system but without liposomes), without any fluorescent sample. The donor signal (Exc 532 nm; Em 593 nm), the FRET signal (Exc 532 nm; Em 684 nm) and the acceptor signal (Exc 638 nm; Em 684 nm) are shown. **(C)**Scheme of smFRET with BmrA labelled with sCy3 and sCy5 at C436, reconstituted in a liposome immobilized with neutravidin on a coverslip passivated with PEG/PEG-biotin. **(D)**Control for the specificity of the liposomes tethering. Fluorescence image of 0.5% DHPE-Texas Red® liposomes incubated in the chamber for 5 min on a surface (left) treated with 20 µg/mL of neutravidin and (right) not treated with neutravidin. Scale bars: 10 µm. **(E)**Typical TIRF-microscopy images showing the Donor channel (Exc 532 nm Em 593 nm), the FRET channel (Exc 532 nm Em 684 nm), and the merge (Donor: green and FRET: red) **(F)**Control of the integrity of the liposomes after immobilization on the coverslip. Left: Liposomes containing 0.5% DHPE-Texas Red® lipids and the soluble dye pyranine in their lumen are incubated for 5min in the chamber. Potential rupture of the liposomes due to the adhesion via the PEG-biotin neutravidin interaction is detected via the fluorescence signal: intact liposomes correspond to colocalization of the green and red signals. Right: fraction of intact liposomes (per image) as a function of the PEG-biotin density on the surface, 10, 2.5, 1 and 0%. The optimal condition is obtained at 1% (*), which has been used in the whole study. 21 images were analyzed. Scale bars: 10 µm.

**Fig. S7.**
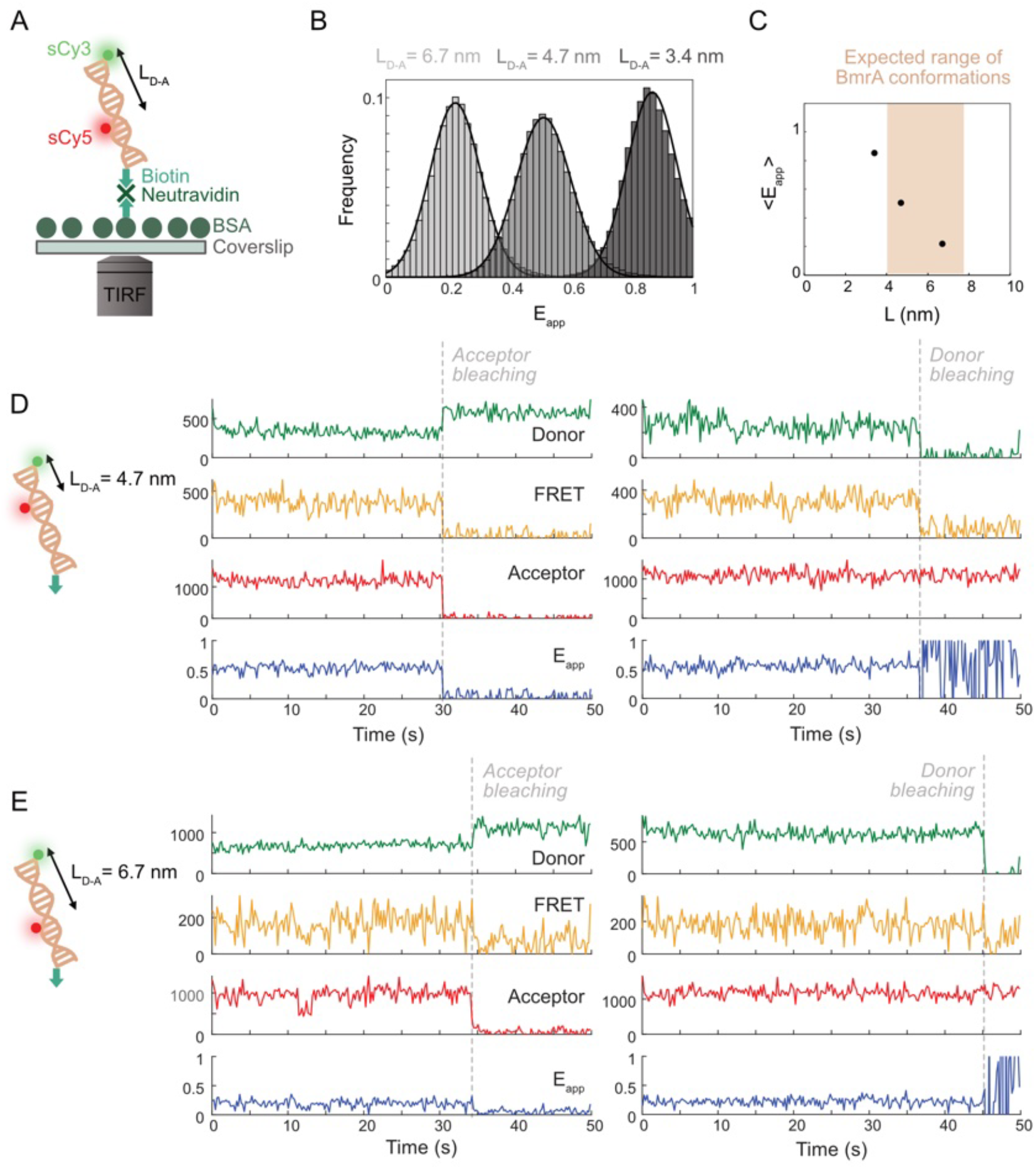
smFRET calibration using a DNA ruler system. **(A)** Scheme of the assay using DNA. 40 base-pair (bp) double-stranded DNAs are labelled with sCy3 (Donor D) and Cy5 (Acceptor A) at different positions for sCy5 (19. 12 and 7 bps), corresponding to the distances between the dyes: L_D-A_ = 6.7, 4.7 and 3.4 nm, respectively. **(B)** Distributions of the apparent FRET efficiency E_app_ for the 3 DNA samples (N = 443, 460 and 538 for 19, 12 and 7 bps, resulting more than 10^5^ time points). **(C)** Mean apparent FRET efficiency <E_app_> deduced from the Gaussian fit of the respective histograms (thick lines in B) as a function of the distance between the dyes (Supplementary Table S1). **(D-E)** Typical intensity signals of donor (green), FRET (yellow), direct acceptor (red) and corresponding apparent FRET efficiency E_app_ (blue) according to time for an interdye distance (D) L_D-A_ = 4.7 nm and (E) L_D-A_ = 6.7 nm. Dashed grey line indicates the acceptor or donor bleaching time. Experiments were performed at 20^°^C.

**Fig. S8.**
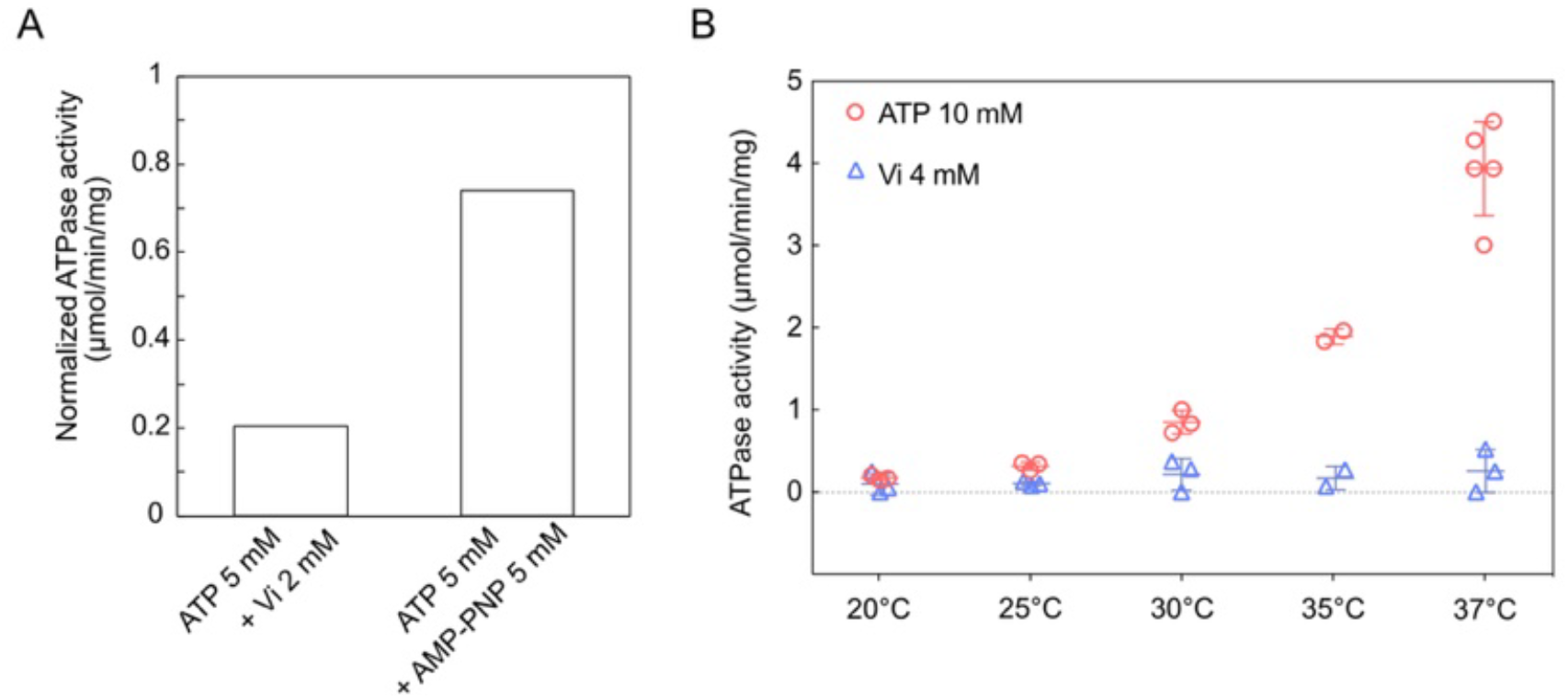
Protein activity in the presence of ATP analogs. **(A**) Normalized ATPase activity measured by spectrometry in vesicles (LV). In the presence of 5 mM ATP, BmrA ATPase activity is reduced by 80% with 2 mM Vanadate (well above K_m_=0.05 mM), but only by 25% with 5 mM AMP-PNP. Note that there is no ATP in our smFRET experiments with AMP-PNP (Fig. 3). **(B)** Protein ATPase activity as a function of temperature, measured by spectrometry in detergent (Anapoe® X-100 at 0.05%). We have checked that the protein remains inhibited by 4 mM Vanadate, independently of the temperature (blue triangles) whereas it changes significantly upon addition of 10 mM ATP (red circles). (Between 2 and 5 measurements).

**Fig. S9.**
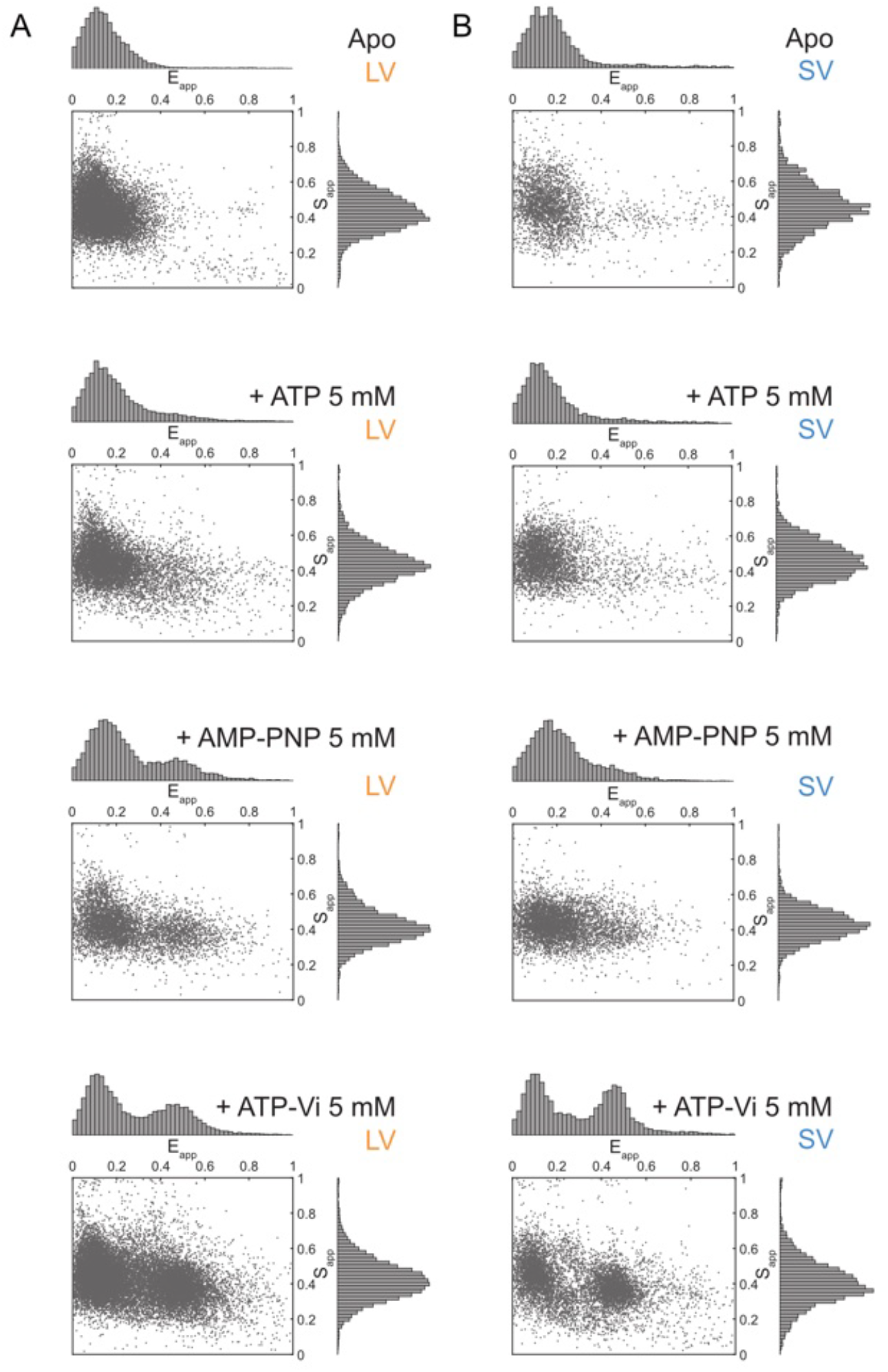
2D smFRET histograms efficiency versus stoichiometry in all conditions at RT (T=20^°^C). Two-dimension histograms of the FRET apparent efficiency E_app_ versus the apparent stoichiometry S_app_. All instantaneous values are depicted. From top to bottom, the protein is in Apo, ATP 5 mM, AMPPNP 5 mM, ATP Vanadate 5 mM, respectively. We display data after selection process, i.e. Donor-Acceptor pairs before photobleaching of a dye. **(A)** Histograms for LV with the total numbers of data points (from top to bottom): N_all_ = 10451, 7743, 5109 and 18559. **(B)** Histograms for SV with N_all_ = 2797, 3959, 5094 and 7552.

**Fig. S10.**
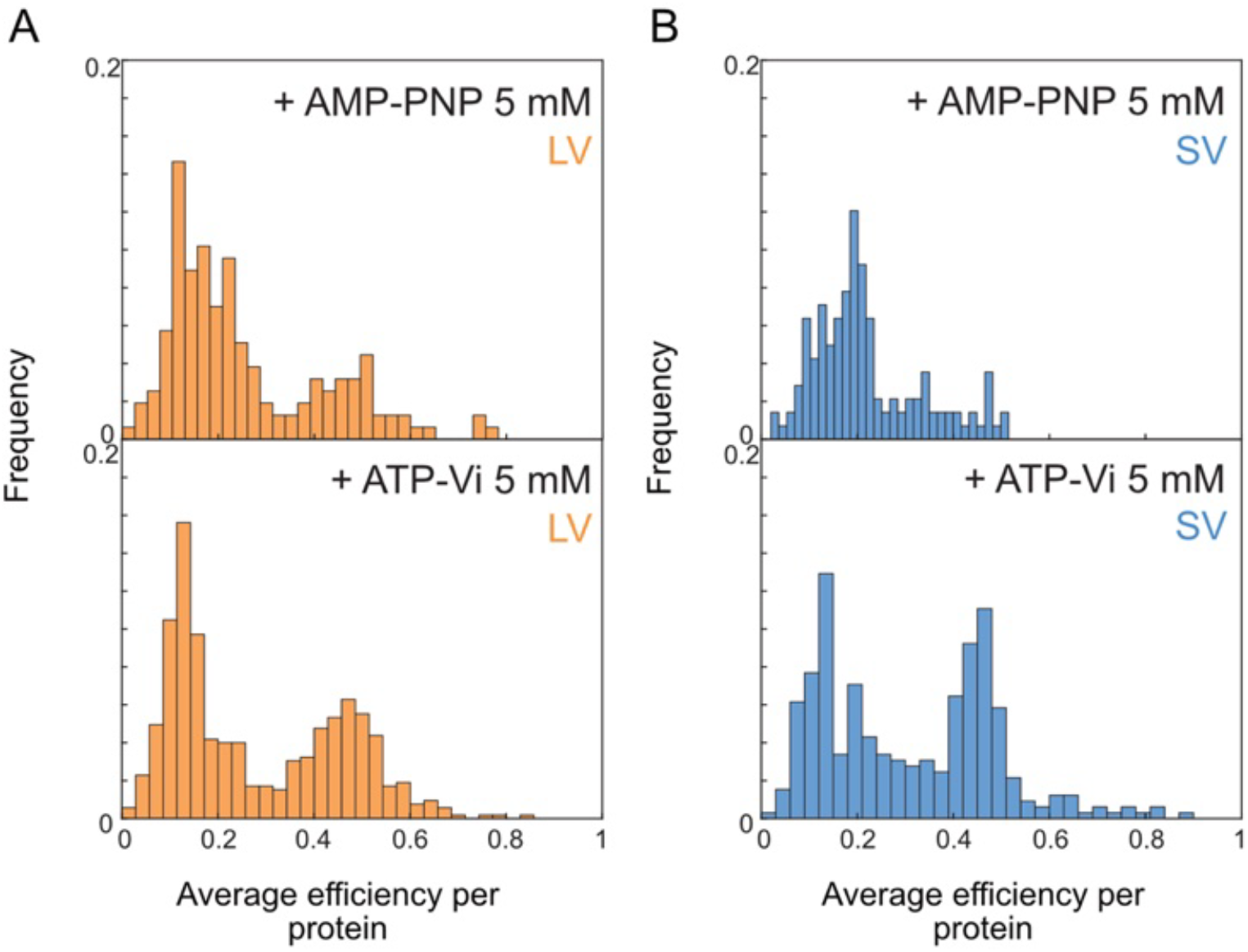
Time average FRET efficiency E_avg_ per protein. Normalized histograms of the FRET efficiency averaged over time for each protein with AMP-PNP 5 mM (top) and ATP Vanadate 5 mM (bottom) in **(A)** LV (orange) N_prot_ = 157 (top) and 525 (bottom) and **(B)** SV (blue) N_prot_ = 141 and 325.

**Fig. S11.**
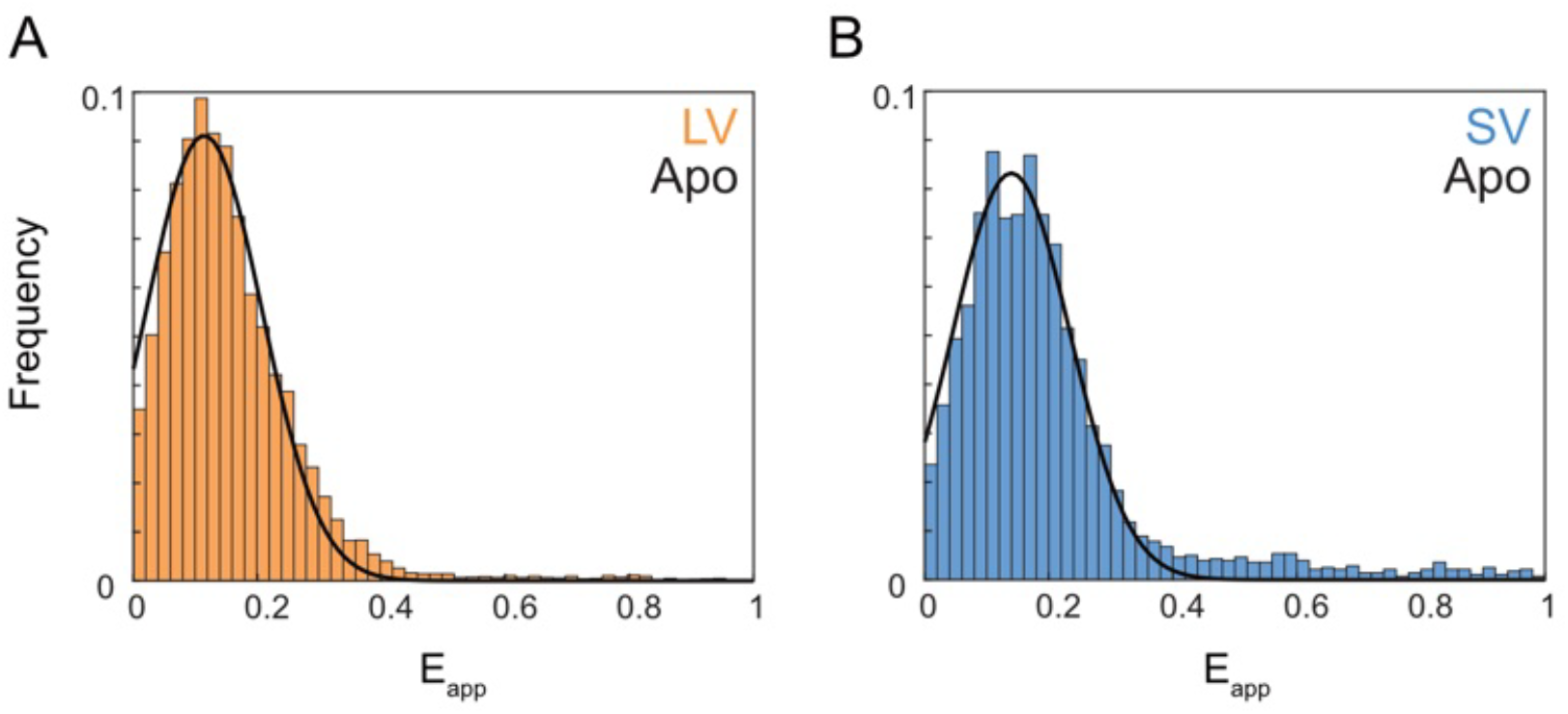
Single Gaussian fit on apo-BmrA FRET distribution. Cumulated histogram of apparent FRET efficiency E_app_ for: **(A)** Apo-BmrA in V125 (N_all_=10451 and N_prot_=310). A single Gaussian fit (thick line) gives: amplitude a = 0.091, mean FRET efficiency <E_app_> = 0.11 and width w = 0.13. **(B)** Apo-BmrA in V20 (N_all_=2797 and N_prot_= 115). The single Gaussian fit (thick line) gives a = 0.083, <E_app_> = 0.14 and w = 0.13.

**Fig. S12:**
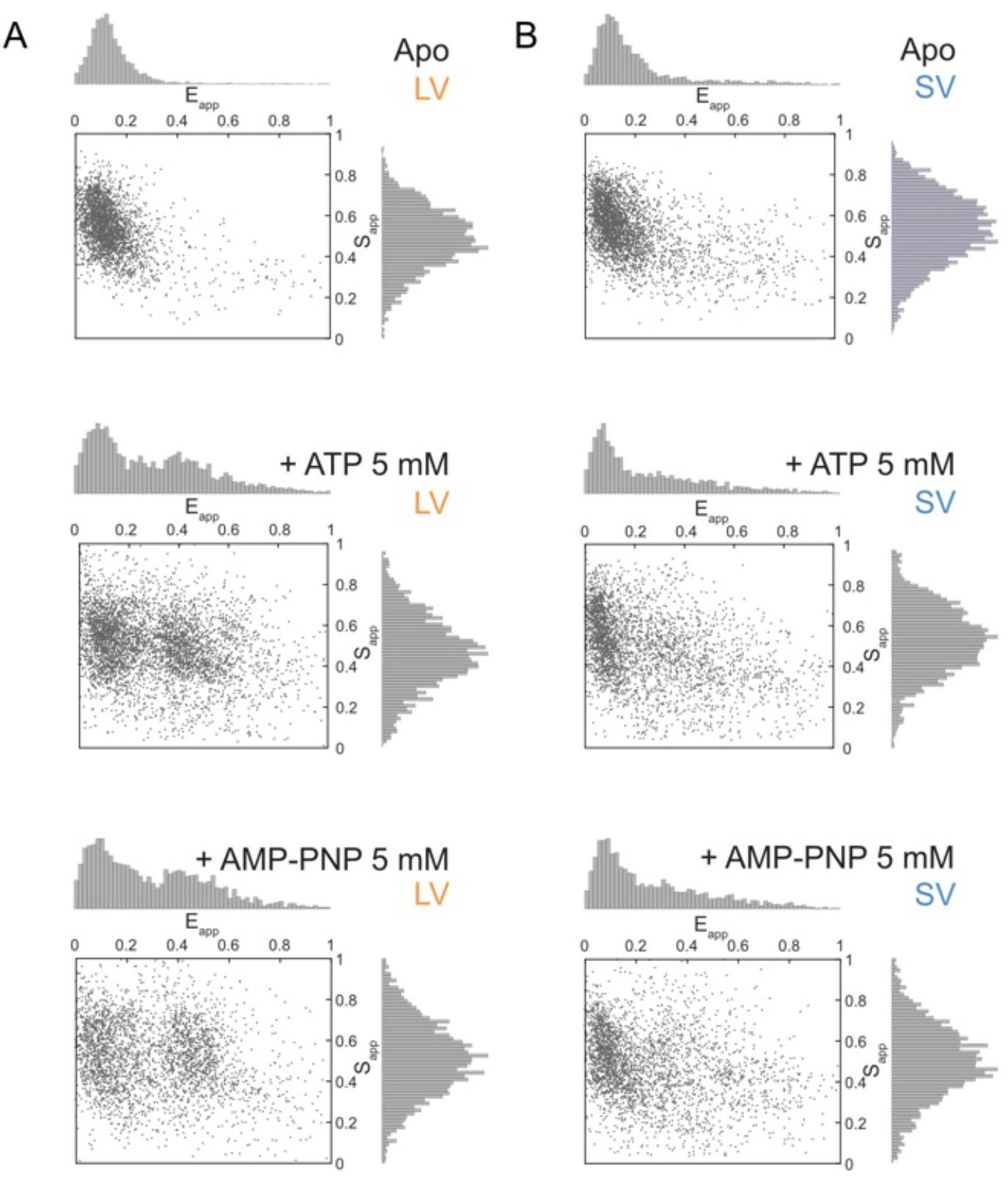
2D smFRET histograms efficiency versus stoichiometry in all conditions at HT (T=33^°^C). Two-dimension histograms of the FRET apparent efficiency Eapp versus the apparent stoichiometry Sapp. All instantaneous values are depicted. From top to bottom, the protein is in Apo, ATP 5 mM, AMPPNP 5 mM, respectively. We display data after selection process, i.e. Donor-Acceptor pairs before photobleaching of a dye. **(A)** Histograms for LV with the total numbers of data points (from top to bottom): N_all_ = 2726, 2991 and 2759. **(B)** Histograms for SV with N_all_ = 2289, 3235 and 2935.

**Fig. S13:**
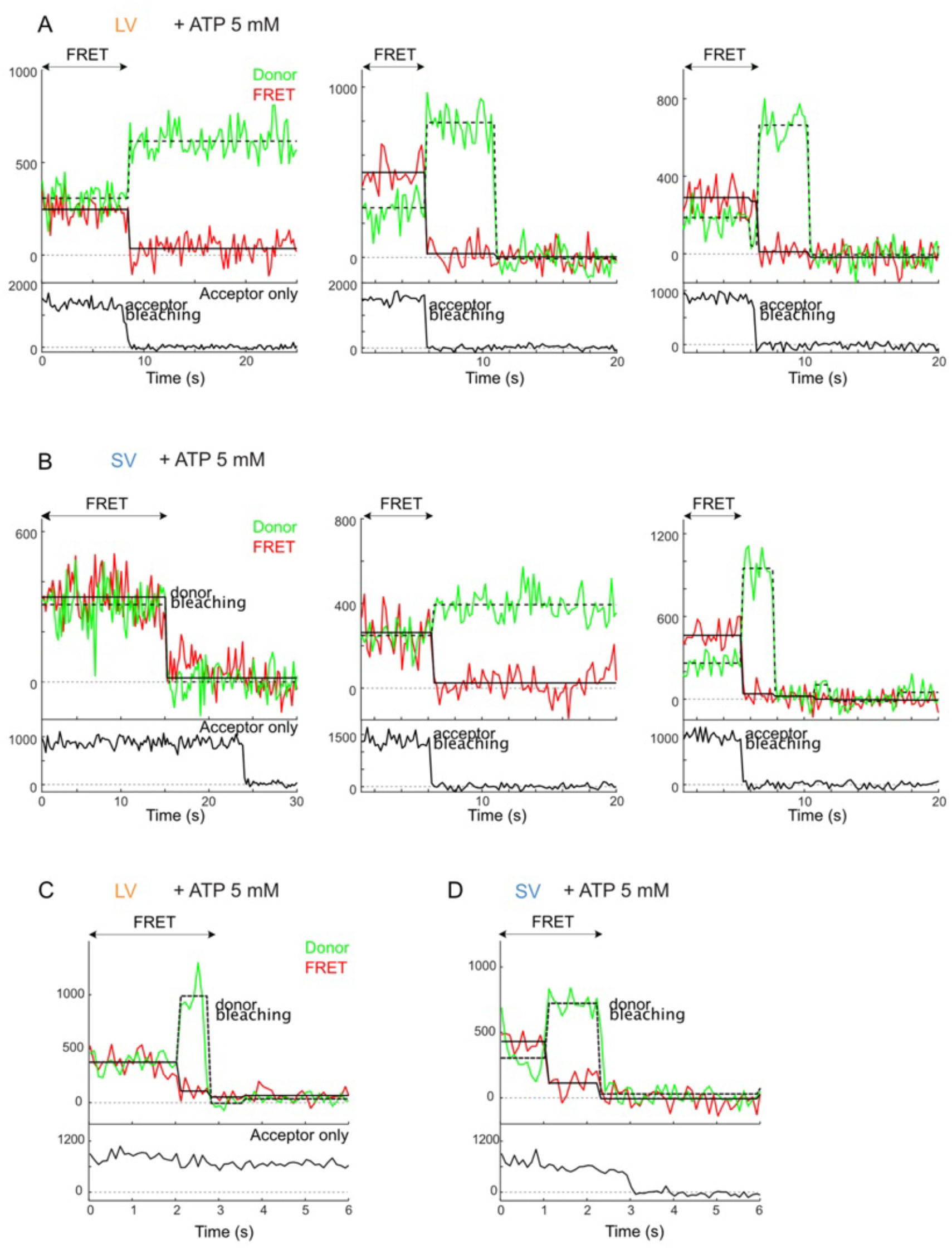
Dynamics of ATP-induced conformational changes at RT (20^°^C). smFRET intensity time traces collected in LV (A-C) and SV (B-D), upon addition of ATP. **(A-B)** On three examples, a stable high FRET signal is measured until either the donor or acceptor dye bleaches, correlating with a drop of FRET signal. Plain and dashed line represent a fit of the FRET switches. **(C-D)** A conformational change is detected with anti-correlated donor (green) and FRET (red) signal, when acceptor intensity (black) remains constant, demonstrating a dynamical event. These switches were observed only twice over 226 proteins for the LV and 3 times over 130 proteins for the SV.

**Fig. S14:**
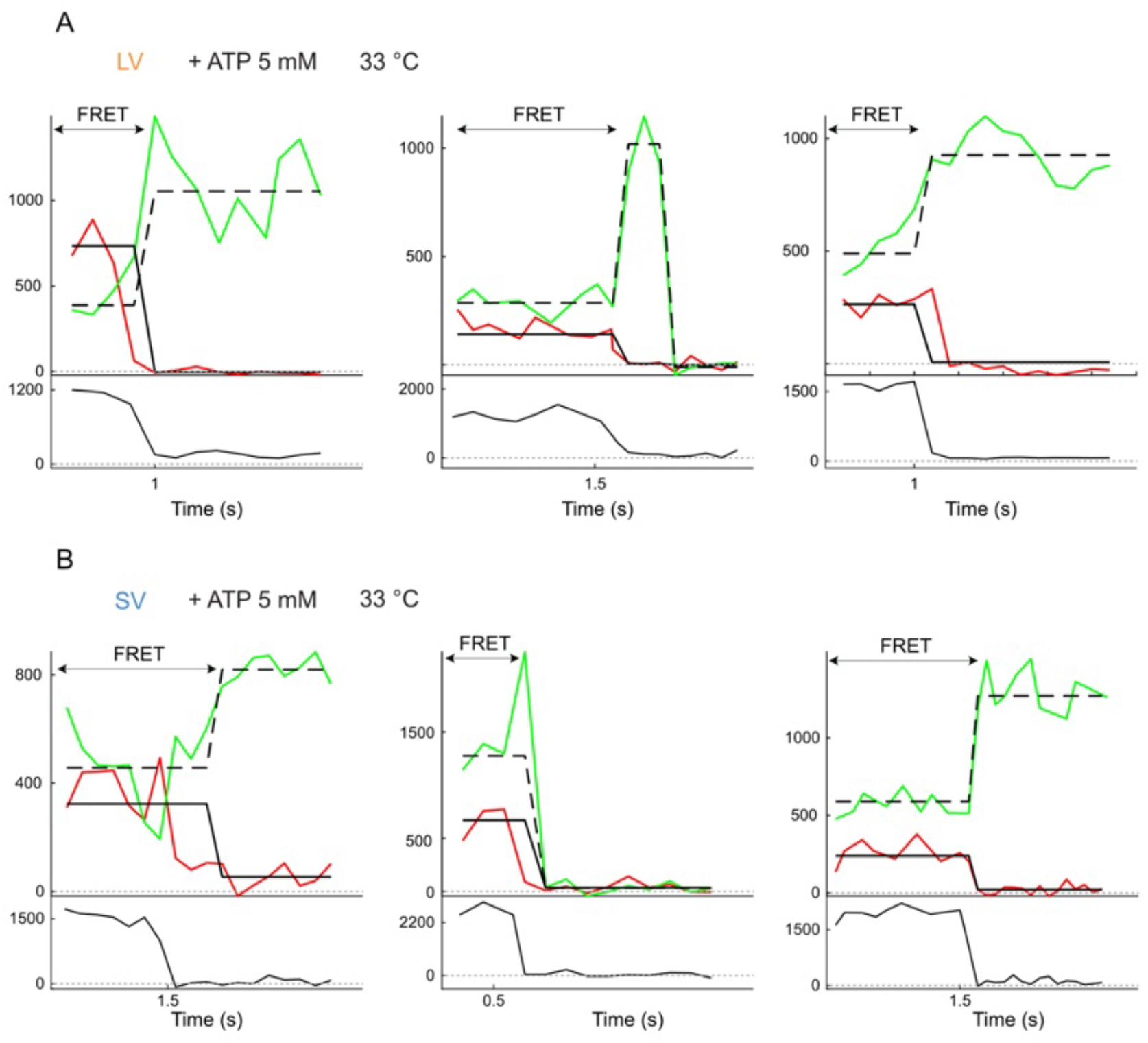
Dynamics of ATP-induced conformational changes at HT (33^°^C). smFRET intensity time traces collected in LV (A) and SV (B), upon addition of ATP. The FRET disappears rapidly (after 1 to 1.5 s) due to photobleaching. No clear conformational switch is detected over this time period.

**Fig. S15:**
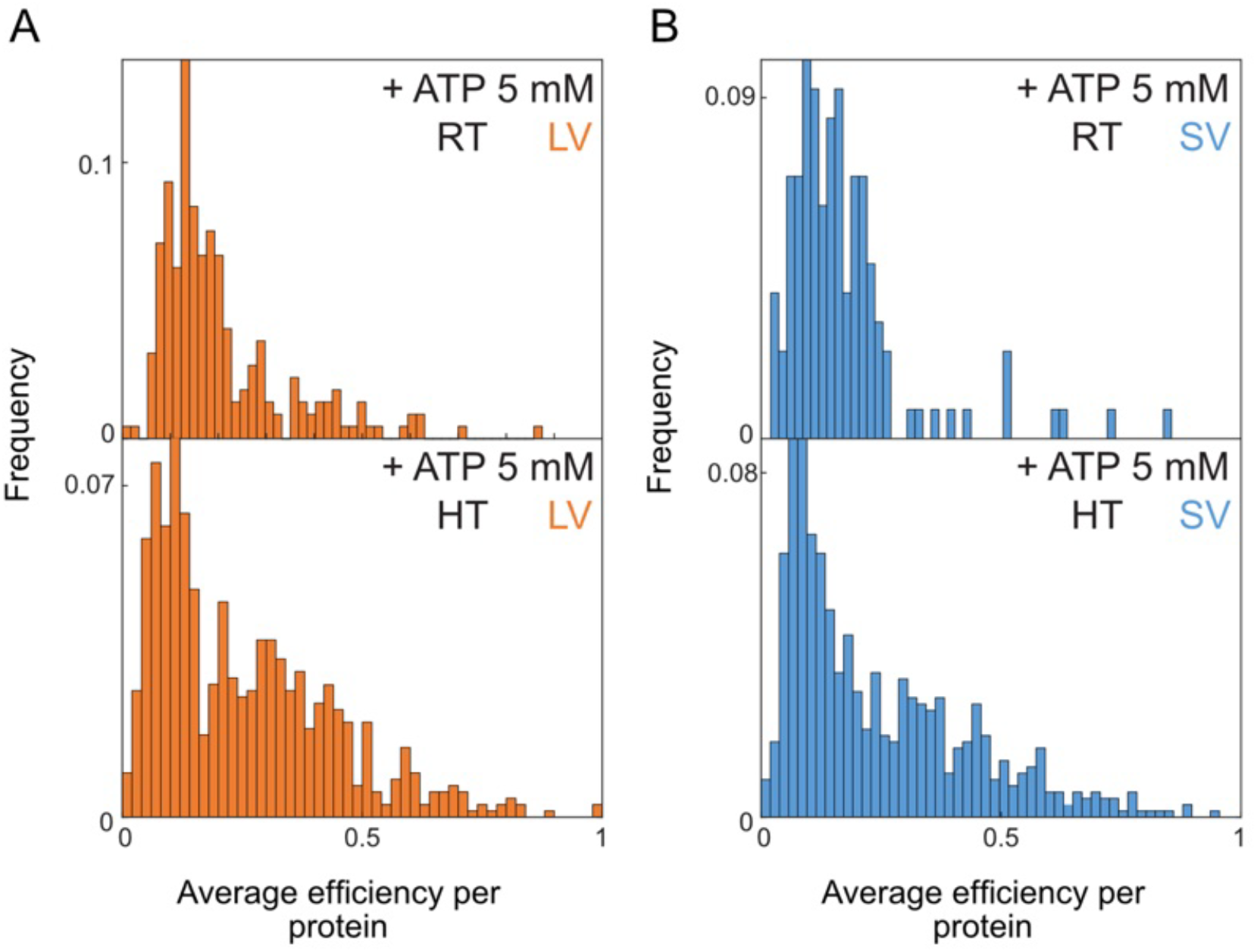
Time average FRET efficiency E_avg_ per protein at RT (20^°^C). Normalized histograms of the FRET efficiency averaged over time for each protein with 5 mM ATP (A) in LV (orange) N_prot_ = 226 (top) and N_prot_ = 748 (bottom) and (B) in SV (blue) N_prot_ = 130 (top) and N_prot_ = 685 (bottom).

**Fig. S16.**
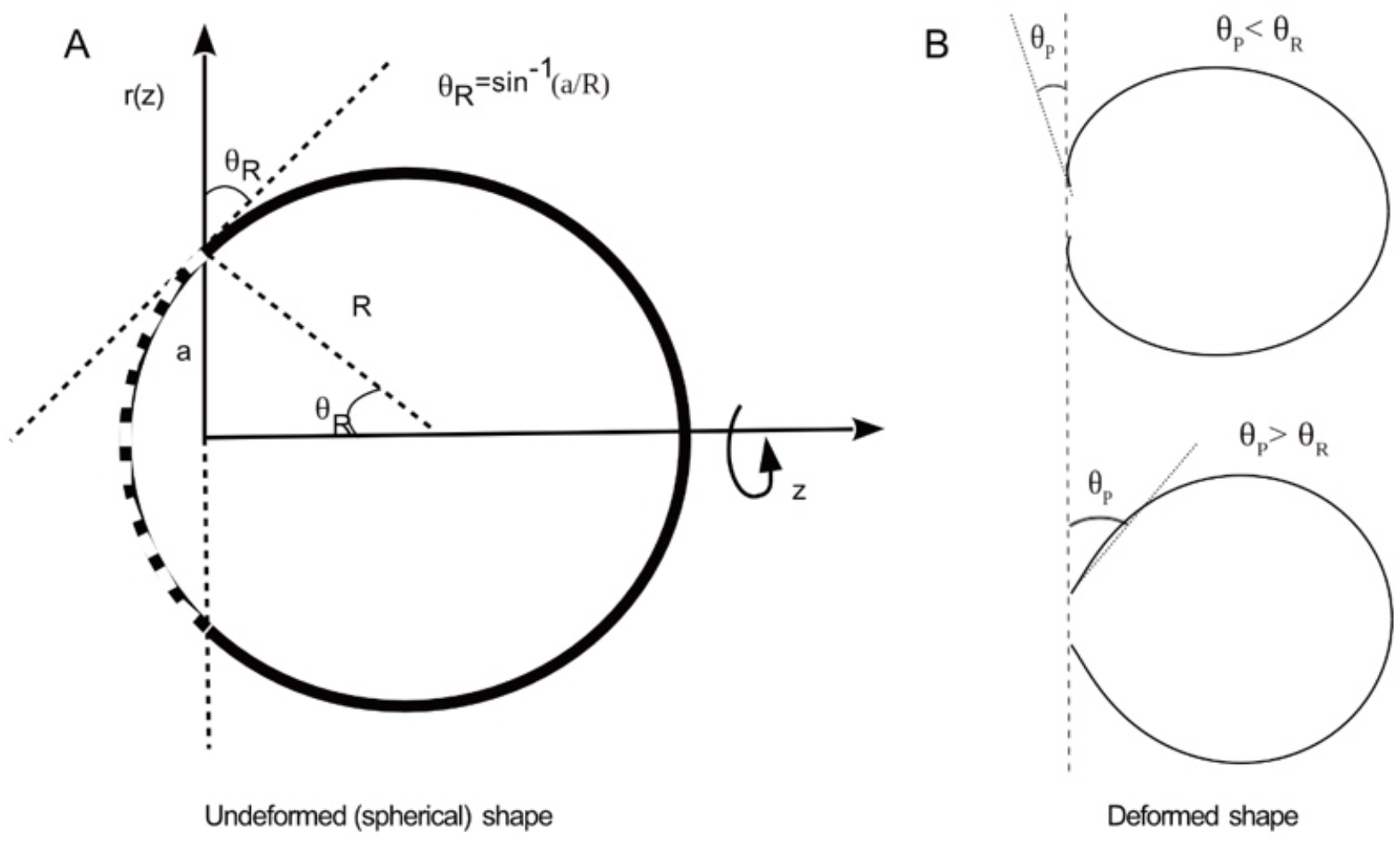
Schematic representation of the model. We consider a vesicle with total area *A*_*0*_ (shown by solid black line in A), having a circular hole of radius *a* (shown by dashed line in A). The full 3D shape of the vesicle can be obtained by rotating the arc through the axis of symmetry *z*. The vesicle can take (A) a spherical shape or (B) a deformed shape depending upon the opening angle *θ*_*P*_.

**Fig. S17:**
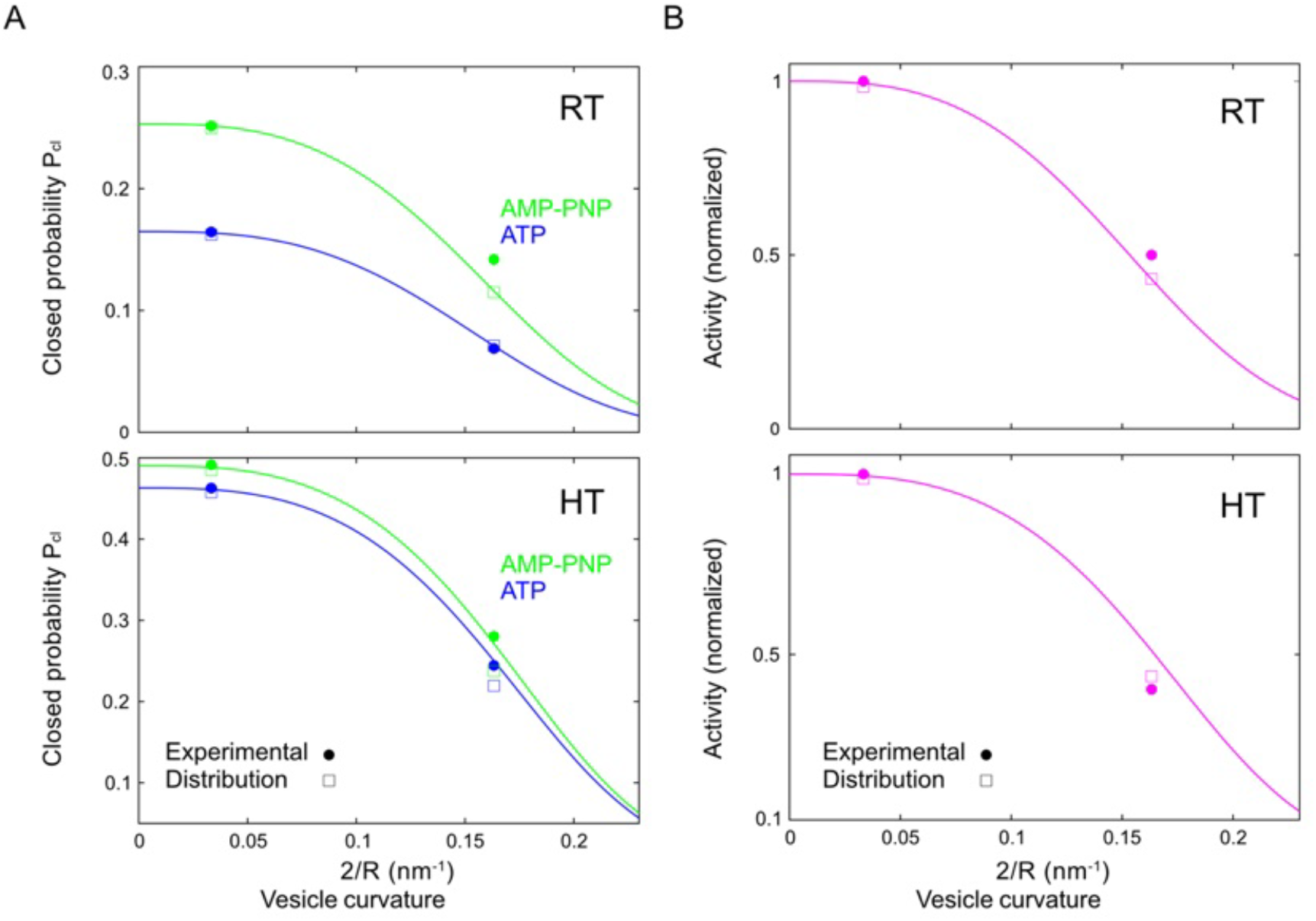
Comparison with experimental results and effect of size distribution on the model. **(A**,**B)** At room temperature (RT). **(C**,**D)** At high temperature (HT). **(A, C)** Fraction of closed population *P*_*cl*_ versus liposome curvature, for ATP (blue) and AMP-PNP (green). The lines are the analytical calculations, circles are the experimental data and the open boxes are the data points that are calculated analytically using the full size distribution of the liposome. **(B, D)** Activity normalized to the activity in a flat membrane versus curvature. Other parameters are the same as in Fig. 4, main text. We fit the experimental data with 2*a*=4nm, *θ*_*op*_=π/10, *θ*_*cl*_=-π/10, and obtain *κ*=27.5 k_B_T.

**Fig. S18:**
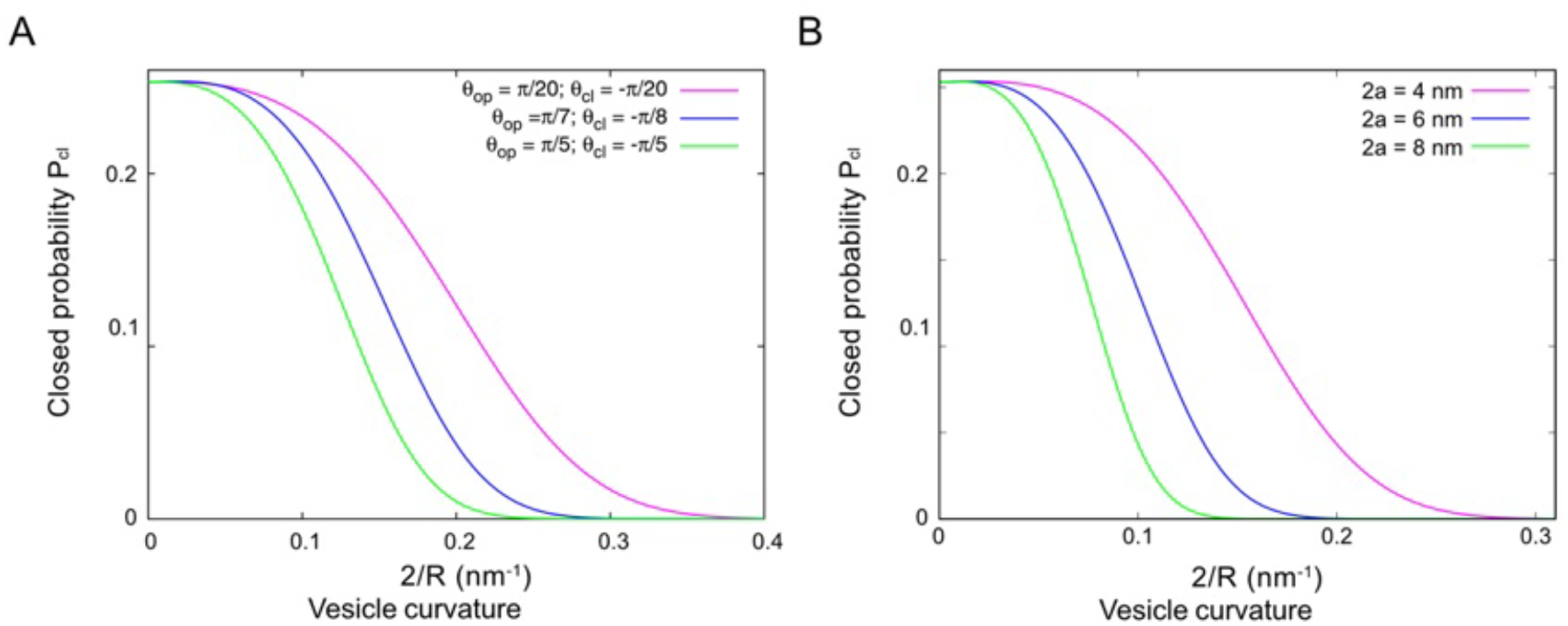
Effect of protein shape and size on the fraction of closed population. **(A)** Calculated fraction of closed population *P*_*cl*_ versus liposome curvature for various values of *θ*_*op*_ and *θ*_*cl*_ (2*a*=4 nm). For higher difference in *θ*_*op*_ and *θ*_*cl*_, the curve is sharper. **(B)** Fraction of closed population *P*_*cl*_ for different protein sizes 2*a* (*θ*_*op*_ =-θ_cl_ =π/10). For larger protein size, the fraction *P*_*cl*_ drops faster with curvature. For both the plots, we use κ=27.5 k_B_T.

**Table S1.** BmrA activity (in µmol ATP/min/mg) corrected by the inward facing fractions of proteins in the different conditions (determined as in Fig. 1C, main text)

| Condition |  | Activity | Corrected activity |
| --- | --- | --- | --- |
| 20 °C | SV | 0.130 | 0.150 |
|  | LV | 0.260 | 0.310 |
| 33 °C | SV | 1.007 | 1.208 |
|  | LV | 2.265 | 2.808 |

**Table S2.**
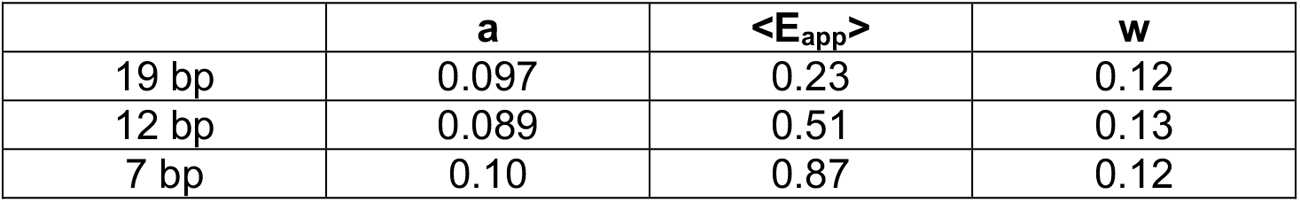
One-gaussian fit parameters of the smFRET histograms obtained for DNA calibration: amplitude *a*, mean *<E*_*app*_*>* and width parameter *w*.

## Supplementary Information for Conformational Changes of the ABC Transporter BmrA Depend on Membrane Curvature

In this document, we derive the energetic cost associated to the membrane deformation upon conformation changes of a membrane protein embedded in a liposome of radius *R*. The membrane energy is related to the local membrane curvature *H* through the Helfrich energy

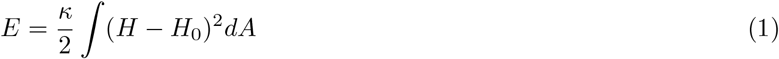

where κ is the bending rigidity, *H*_0_ the spontaneous curvature and the integral is over the liposome surface *A*. The liposome is assumed to have an optimal conformation in the absence of protein, so that the spontaneous curvature is related to the vesicle radius according to *H*_0_ = 2*/R* and the optimal shape is a sphere of radius *R*.

Here we assume that the protein is an axisymmetric cone of radius *a* and opening angle *θ*_*P*_ (defined positive when facing outward). We assume that only the opening angle varies upon shape transformation while the protein radius remains unchanged. Consequently, the optimal liposome shape remains axisymmetric upon protein conformation change. The different geometric quantities are defined in Fig.S16.

### I. SMALL DEFORMATION

Here we concentrate on small deviations around a sphere, which requires that the angle imposed by the protein *θ*_*P*_ is close to the angle that would perfectly fit the membrane natural shape, namely *θ*_*R*_ = *sin*^−1^(*a/R*).

Defining the local distance of a membrane segment to the axis *r*(*s*) and the local angle of the membrane normal with the axis *θ* (*s*) (*s* is the curvilinear coordinate, see Fig.S16(A)), we have:

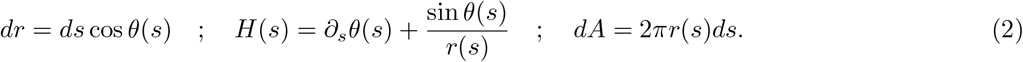

The non-deformed spherical shape satisfies *θ*_0_(*s*) = *θ*_*R*_ + *s/R* and *r*_0_(*s*) = *R* sin *θ*_0_(*s*). Assuming small variations of the membrane shape *r*(*s*) = *r*_0_(*s*)+ Δ*r*(*s*), we have

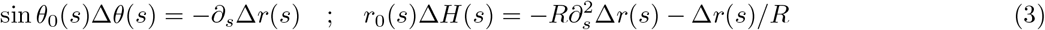

The energy variation associated to a change of shape reads at lowest order

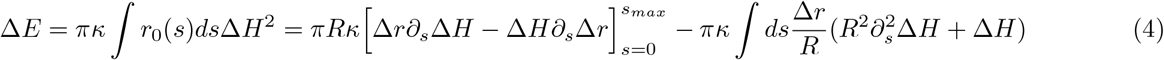

where the RHS of the equation has been obtained after two integrations by part. Minimisation of the energy difference Δ*E* leads to the Euler Lagrange equation,

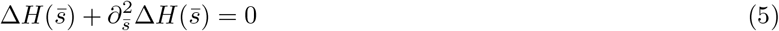

where, we use 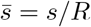. The curvature and abscissa variation can be written as,

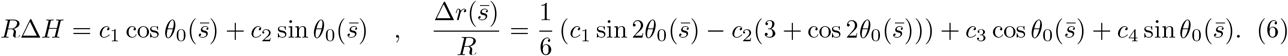

The integration constants *c*_1_, *c*_2_, *c*_3_, *c*_4_ must be determined from the boundary conditions (BCs)

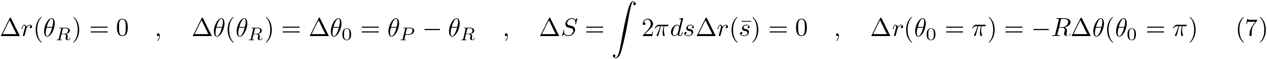

The first BC indicates that the protein areal footprint does not change, the second that the protein opening angle changes by a value Δ*θ*_0_ = *θ*_*P*_ − *θ*_*R*_, the third BC indicates that the total liposome area does not change, and the last enforces that 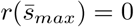 and 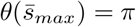 at the liposome pole opposite to the protein.

The set of four equations for the four integration constants can be solved for arbitrary ratio *a/R*. Here we focus on situations where the protein is much smaller than the liposome (*a* ≪ *R*), for which we obtain:

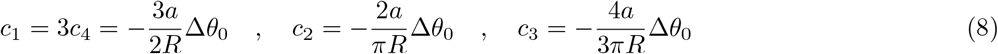

From this the energy cost of the change of angle can be obtained from Eq.4:

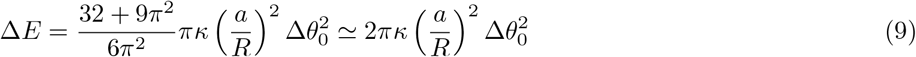

The energy of the membrane can thus be written as,

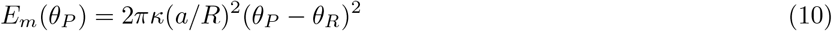

Thus, the total energy of the system can be written as,

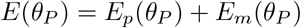

where, *E*_*p*_ is the energy of the protein. The energy difference between the protein in the closed state and in the open state is written as

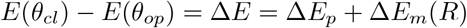

With

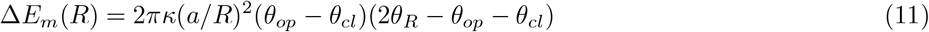

In the following, we assume that the intrinsic protein energy *E*_*p*_ is independent of the liposome radius, which only influences the membrane energy *E*_*m*_.

### II. STATISTICAL ANALYSIS OF THE LIPOSOME POPULATION, AND COMPARISON WITH EXPERIMENTAL DATA

If *N*_*op*_ is the population of the liposomes in the open conformation (low FRET), *N*_*cl*_ is the population in the closed conformation (high FRET), *k*_1_ is the rate of transition from ‘open’ to ‘closed’ state, and *k*_2_ is the transition rate from ‘closed’ to ‘open’ state, then the kinetic equation for two probabilities are:

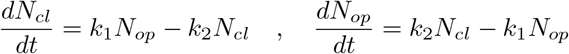

At steady state, both populations will be constant in time, i.e., 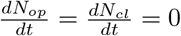, which gives,

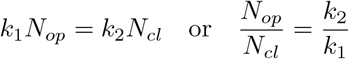

We may assume that the ‘open’ to ‘closed’ transition is driven by thermal fluctuations. In experiments, ATP and its different analogues have been used, among which we are interested in mainly two cases: (1) Non-hydrolizable ATP (AMP-PNP), which is an equilibrium case, where the closed state is stabilized by a binding energy ϵ between two nucleotides and thermal fluctuations can lead to unbinding and allow the transition from the closed to the open state; (2) Hydrolizable ATP (ATP), where ATP hydrolysis is required to destabilise the closed state and allow the transition to the open state.

For the equilibrium case (AMP-PNP), the trasition rates are given by the Arrhenius law: 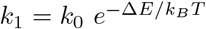, and 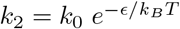, where *k*_0_ is a constant having unit of inverse time. We assume that the energy barrier associated to membrane deformation is the energy difference between the open and closed state, given by Eq. 11. The fraction of open (low FRET) to closed (high FRET) population is then given by,

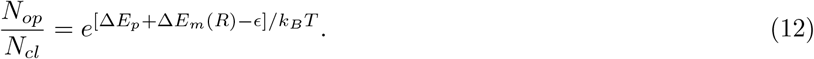

Similarly, for the non-equilibrium case (ATP), assuming *k*_2_ = *k*_*ATP*_ the ATP hydrolysis rate, we have,

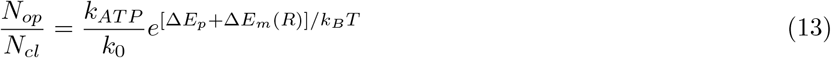

Experimentally, we know the open and closed population for two sets of vesicle sizes: small vesicles (*SV*) and large ones (*LV*). Taking the ratio of the above fraction for two different liposome sizes, we get,

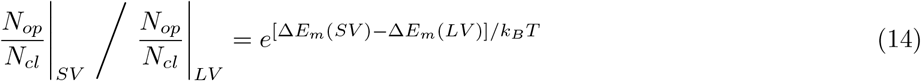

which depends only on the membrane parameter (e.g., bending rigidity *ω*) and is the same for both the equilibrium as well as the non-equilibrium cases.

The experimental data are given in Table 2, main text. In this data, we recalculate the populations by subtracting the high FRET Apo populations from each case, that are considered to be non-functional. Using the above ratio (Eq. 14), we can calculate the value of *ω* for both the AMP-PNP and ATP cases. For room temperature (RT), the calculation gives us the value of κ ≃ 21.0*k*_*B*_*T* for the AMP-PNP case and κ ≃ 29.5*k*_*B*_*T* for the ATP case. Similarly, for high temperature (HT), the calculation gives us the value of κ ≃ 26.0*k*_*B*_*T* for the AMP-PNP case and κ ≃28.5*k*_*B*_*T* for the ATP case. These values are rather close, and allows us to fit the experimental data for a moderate κ (both for RT and HT), that we discuss below.

For the equilibrium case, the fraction of population in the closed state is given by,

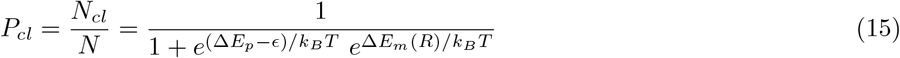

where, *N* = *N*_*cl*_ + *N*_*op*_ is the total population. We can verify that for LV case, the membrane contribution is negligible as it is much smaller than *k*_*B*_*T*, such that we can write 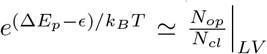. We can use this value in the above equation, and plot *P*_*cl*_ as a function of the liposome curvature. In the experiment, the liposome sizes are measured by considering its outer surface, thereby we subtract the bilayer thickness (∼ 5*nm*) from the actual data set of diameter when we compare with the analytical results. We show this plot in Fig. S17(A), green line for RT. The circles are the experimental data points that fits well for κ = 27.5 *k*_*B*_*T*. Other parameters we use here are: 2*a* = 4 *nm, θ*_*op*_ = *θ/*10 and *θ*_*cl*_ = −*π /*10. The open boxes are the data points that are calculated analytically using the full distribution of liposome size,

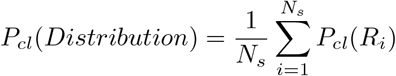

We perform similar calculations for the non-equilibrium case, which gives,

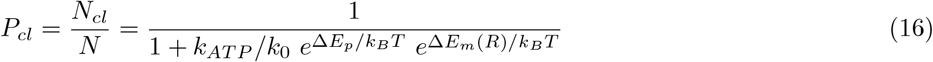

In a similar way as above, we can approximate 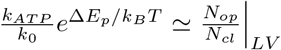, and use it to the above expression. We show this plot in Fig. S17(A), blue line. Here also, we compare with the experimental data as well as the data points calculated using the full distribution of liposome size.

We also compare the activity of the ATP for RT, which is defined as the rate of ATP hydrolysis per mole. We define the activity as,

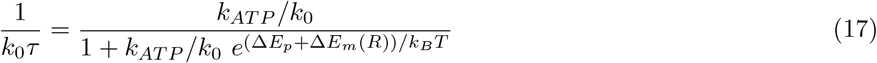

We plot the normalized activity in Fig. S17(B) for RT. We note that ATP activity decreases with the curvature of the liposome, but the effect is strong only for higher curvature.

Similar calculations are also performed for HT and shown in Fig. S17(C-D) for the same value of fitting parameter κ = 27.5*k*_*B*_*T*.

### III. EFFECT OF PROTEIN CONFORMATION AND SIZE ON THE FRACTION OF CLOSED POPULATION

In the above discussion, we use the parameter values that are applicable for BmrA protein. Here, we would like to discuss how the results are affected when we consider different values of *θ*_*op*_ and *θ*_*cl*_, and also when we change the protein size *a*. In Fig. S18(A), we plot the fraction of population in the closed state as a function of curvature, which shows that larger difference in *θ*_*op*_ and *θ*_*cl*_ makes the curve sharper. The effect of protein size is also prominent (Fig. S18(B)), and for a protein which is twice the BmrA, *P*_*op*_ shows sharp increase with curvature, even in the smaller curvature regime. This emphasizes the fact that the effect of membrane mechanics on the transport process may be very effective for other kind of proteins with larger size, such as piezo1.

